# NAD^+^ - and EVA1-C-dependent reversal of neurological deficits is mediated by differential alternative RNA splicing in tauopathic animal models

**DOI:** 10.1101/2024.10.27.620478

**Authors:** Ruixue Ai, Lipeng Mao, Xurui Jin, Shi-qi Zhang, Junping Pan, Maria Jose Donate Lagartos, Shu-Qin Cao, Guang Yang, Chenglong Xie, Xiongbin Kang, Pingjie Wang, Yang Hu, Linda Hildegard Bergersen, Jon Storm-Mathisen, Hidehito Kuroyanagi, Beatriz Escobar Doncel, Noemí Villaseca González, Farrukh Abbas Chaudhry, Zeyuan Wang, Qiang Zhang, Zhangming Niu, Guobing Chen, Oscar Junhong Luo, Evandro Fei Fang

**Affiliations:** Department of Clinical Molecular Biology, University of Oslo and Akershus University Hospital, 1478 Lørenskog, Norway; Guangdong-Hong Kong-Macau Great Bay Area Geroscience Joint Laboratory, 510632 Guangzhou, China; Department of Systems Biomedical Sciences, School of Medicine, Jinan University, 510632 Guangzhou, China; Mindrank AI Ltd, Hangzhou, China; Department of Microbiology and Immunology, School of Medicine; Institute of Geriatric Immunology, School of Medicine, Jinan University, 510632 Guangzhou, China; Cardiovascular Research Centre, Royal Brompton Hospital, SW7 2AZ London, United Kingdom; National Heart and Lung Institute, Imperial College London, SW7 2AZ London, United Kingdom; Department of Neurology, the First Affiliated Hospital of Wenzhou Medical University, Wenzhou, 325000 Zhejiang, China; Institute Of Aging, Wenzhou Medical University, Wenzhou, 325000 Zhejiang, China; Genome Data Science, Faculty of Technology, Bielefeld University, Bielefeld, Germany; Department of Geriatrics, the First Affiliated Hospital, Zhengzhou University, 450052 Zhengzhou, China; Institute of Oral Biology (IOB), UiO, Faculty of Dentistry, 0372 Oslo, Norway; the Center of Healthy Aging (CEHA), Medical Faculty, University of Copenhagen, 2200, Denmark; Division of Anatomy, Department of Molecular Medicine, Institute of Basic Medical Sciences, University of Oslo, NO-0317 Oslo, Norway; Department of Biochemistry, Graduate School of Medicine, University of the Ryukyus, Okinawa 903-0215, Japan; Faculty of Pharmacy and Regional Centre of Biomedical Research (CRIB), University of Castilla-La Mancha, 02008 Albacete, Spain; Department of Molecular Medicine, University of Oslo (UiO), 0372 Oslo, Norway; College of Computer Science and Technology, Zhejiang University, Hangzhou 310027, China; Hangzhou Global Scientific and Technological Innovation Center, Zhejiang University, Hangzhou 311200, China; The Norwegian Centre on Healthy Ageing (NO-Age), 1478 Oslo, Norway

## Abstract

Aberrant alternative splicing (ASEs) is an aging hallmark to Alzheimer’s Disease (AD). Although NAD^+^ and related metabolites can slow down AD, NAD^+^ on ASEs in AD remain unclear. Mouse transcriptomic data revealed NR-induced ASEs, focusing on the Eva1-C locus. AI-based algorithms predicted EVA1-C protein structures and protein-protein interactions. AD postmortem brain samples and tauopathy models including transgenic mice and worm was used for validation. NAD^+^ abundance/metabolic status modulates ASEs and the expression of EVA1-C isoforms, which in turn regulate the interaction with BAG-1 and HSP70 proteins. Importantly, EVA1-C is dramatically reduced in 20 Braak 5/6 AD patients compared to cognitive normal humans in different brain regions. NAD^+^ metabolism modulates abundance of specific mRNA isoforms, and that ASEs influence disease progression in model tauopathies and potentially AD. These results could facilitate future development of NAD^+^-based splice-switching therapeutics for AD.

**Teaser:** Unveiling the Link Between NAD^+^ Metabolism and Alzheimer’s Disease: Discovering the Role of Alternative RNA Splicing in Disease Progression and Potential Therapeutic Targets

## Introduction

Dementia is a prevalent and devastating disease that is reported by the World Health Organization to be the 7^th^ leading cause of mortality worldwide (*1*). Alzheimer’s disease (AD) is the most common form of dementia, few effective treatments are currently available for AD, and it is estimated to affect more than 50 million people globally (*1*). Intracellular neurofibrillary tangles (NFTs) composed of hyper-phosphorylated Tau (pTau) are a prominent neuropathological feature of AD brains (*2, 3*). Research on genetic risk factors for AD identified several genes including mutant alleles of human Tau (hTau), which led to the development of valuable hTau-transgenic mouse models for AD (*4–6*).

Alternative splicing events (ASEs) represent an important post-transcriptional regulatory mechanism that affects 92-94% of human genes (*7*). Alternative splicing is unique to eukaryotic cells and is the process of removing introns and assembling exons to construct multiple RNA transcript isoforms from a single pre-mRNA transcript (*8*). The function of one transcript isoform can be related to, distinct from, or even opposite to the function of other transcript isoforms from the same gene (*7, 9, 10*). The processes that regulate alternative splicing are abundant and conserved in the vertebrate nervous system (*11*), resulting in phenotypic differences between individual cells or organisms. Aberrant mRNA splicing events are associated with AD (*12, 13*) and have been documented in candidate AD-associated genes including murine and human microtubule-associated protein tau (*MAPT*) which encoding the Tau proteins (*13, 14*). The exon-3-containing *MAPT* mRNA transcript decreases aggregation of NFTs (*13, 14*). However, analyses of the proteomic profiles of AD patients’ brains identified structural changes in insoluble U1 snRNP, a small nuclear RNA component of the spliceosome (*15*). Furthermore, a global transcriptome association study of 450 subjects in two aging cohorts identified hundreds of ASEs reproducibly associated with AD pathology (*5*). Additional studies support the idea that ASEs are an important feature of AD brains and that ASEs are in some cases modulated by genetic and/or protein factors (*5*).

Emerging evidence suggests that age– and genetic-dependent NAD^+^ depletion and/or impaired NAD^+^-dependent pathways play a role in AD pathophysiology (*16, 17*). In rodent models of early-onset familial AD, NAD^+^ depletion and NAD^+^-associated metabolic dysfunction have been linked to disease pathology in brain tissue (*18*). Meanwhile, injection of an NAD^+^ precursor inhibited Tau phosphorylation and improved cognitive function in a cross-species study involving a neuronal Tau (pro-aggregate F3ΔK280 Tau fragment) transgenic *C. elegans* model (*17*), 3xTgAD/Polβ^+/−^ mice (*19*), and 3xTgAD mice (*20*). However, only a small number of studies have addressed the role of ASEs in tauopathies or other AD-related pathologies (*6*) and a comprehensive mechanistic study of the relationships between NAD^+^ metabolism and ASEs in tauopathies is lacking.

In 2016, the U.S. FDA approved therapeutic use of small molecule compounds and genomics-based technologies including oligonucleotides via direct delivery into the brains of patients afflicted by central nervous system (CNS) diseases (*21*). This type of therapy has been tested preclinically for refractory neurological disorders (*21*) including erythropoietic protoporphyria (*22*), cystic fibrosis (*23*), congenital deafness (*24*), and cancers such as glioblastoma (*25*), prostate (*26*), and breast cancer (*27*). However, the development and application of therapeutics that target mRNA splicing has been challenging, at least in part because the biological effects of ASEs are complex and difficult to predict (*22*). Therefore, there is an urgent need for better understanding of the regulation and downstream effects of ASEs especially in the context of AD, tauopathies and related neurological diseases, with a final goal in developing effective therapies.

Eva1-C plays an important role in neuronal development and function, while its role in AD is obscure(*28, 29*). To understand the biological role of *Eva1-C* and its isoforms in mice with or without a tauopathy phenotype, it is important to identify and understand Eva1-Cs protein-protein interactions (PPIs). Several high-throughput methods for analyzing PPIs are available including yeast two-hybrid screens (Y2H) (*30*), tandem affinity purification (TAP) (*31*), and mass spectrometric protein complex identification (MS-PCI) (*32*); however, these methods are time-consuming and prone to high false-positivity and false-negativity rates (*32, 33*). Recent analyses of PPIs employ novel computer-driven AI-based deep-learning algorithms to predict PPIs. One such approach is DeepPPI, which deduces high-level protein features from protein sequence data and has outperformed traditional machine learning algorithms (*34*). However, currently available methods use protein amino acid sequences (primary structure) as input and therefore may neglect important features of protein secondary and tertiary structures and 3-dimensional (3D) conformations (*35–38*). The present study employs a novel approach, whereby genetic variation, predicted 3D protein structures, and predicted PPIs are modeled using large-scale bioinformatic analyses and deep learning algorithms. The output data are then verified in a “wet”-lab based cross-species experimental system. Using this approach, we performed global transcriptomic analyses and identified ASEs in RNA-seq data from the hippocampi of 14-month-old wild-type (WT) and hTau.P301S mice with or without exposure to NR. Focusing on *Eva1-C*, a deep learning framework was used to predict EVA1-C protein structures and PPIs, its mechanism(s) of action, and EVA1-C isoform-specific binding affinities to BAG-1 and HSP70 in the presence or absence of NR. Importantly, our findings were experimentally verified in mammalian cells, nematodes, mice, and postmortem brain tissues. This comprehensive approach yielded insight into the effects of NR on ASEs in tauopathies and the potential mechanism(s) of action of EVA1-C. The results will inform and improve future design of mRNA splicing-targeted therapeutics and could facilitate successful future clinical use of these therapeutics.

## Results

### Dysregulation of mRNA splicing in transgenic hTau.P301S mice and aged worms

The effects of NR on ASEs and the RNA transcriptome were investigated in hippocampal tissue from 14-month-old hTau.P301S and wild-type (WT) mice (control). Eleven-month-old mutant and WT mice were maintained with or without NR for 3 months, a time period that overlaps the age of expected onset of congnitive impairment and neuronal degeneration in hTau.P301S mice. RNA-seq analyses revealed differential up-or downregulation of 509 genes (FDR<0.05) in hTau.P301S mice relative to WT mice (**Fig. 1a; Table S1**). Gene Ontology (GO) analyses showed that down-regulated genes were enriched especially for RNA processing-related GO biological process and cellular component terms including RNA splicing (richFactor= –0.028), mRNA processing (richFactor= –0.030), regulation of mRNA metabolic process (richFactor= –0.038), regulation of RNA splicing (richFactor= –0.044), and alternative mRNA splicing via spliceosome (richFactor= –0.071) (marked in **Fig. 1b; Table S2**). In contrast, upregulated genes were not enriched at a statistically significant level for GO terms related to RNA splicing (**Table F 1a, Table S2**). This result suggests that dysregulation of mRNA processing, especially RNA splicing, is a hallmark of tauopathy in the humanized hTau.P301S mouse.

**Fig. 1.**
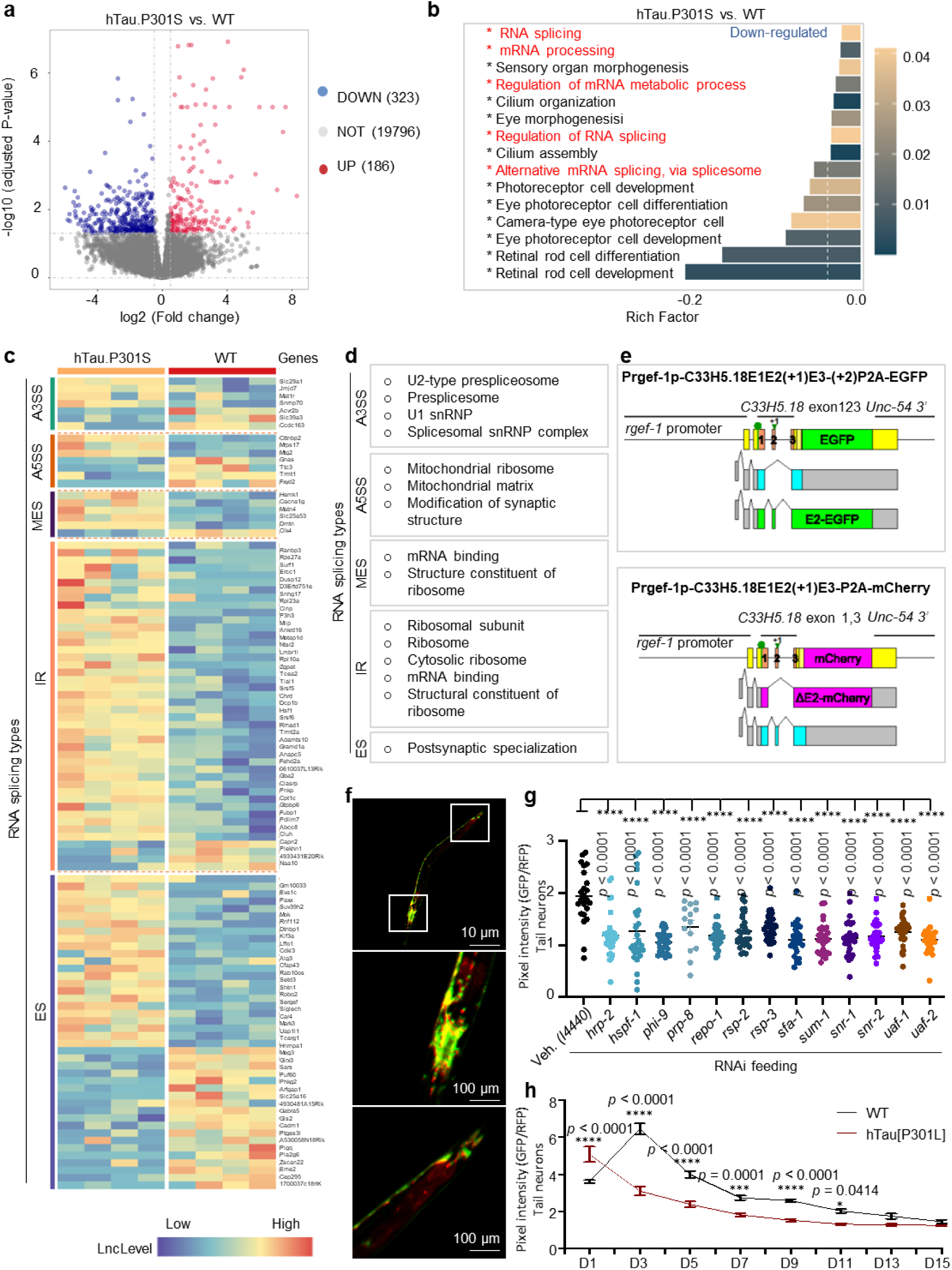
Compromised mRNA splicing in tan pathology and aging. **a.** Volcano plot comparing hippocampal RNA-seq-derived gene expression in hTau.P301S and wild-type (WT) mice. Gray dots represent 19,796 genes expressed at similar levels in both strains; red dots represent differentially upregulated genes and blue dots represent differentially down-regulated genes in mutant relative to WT mice. b. Top 15 GO pathways enriched (adjusted P-value <0.05) among genes down-regulated in hTau,P3OlS vs. WT mice; GO terms shown in red relate to mRNA splicing or processing. The criterion for differential expressed genes (DEGs) is (log2(fold-change) >0.5 and FDR <0.05) c. Heatmaps showing distribution and relative frequency of ASEs stratified by ASE subtype in hTau,P3OlS vs. WT mice; ASE subtypes are: alternative 3’ splice site (A3SS), alternative 5’ splice site (A5SS), multiple exon skip (MES), intron retention (IR), exon skip (ES). d, GO terms enriched in DEGs stratified by ASE subtype, e, Schematic diagram of the *rgef-1* splicing reporter gene cassette, f. Pan­neuronal *rgef-1* splicing in day 1 **C.** *elegans*, **Inset:** positive control reporter cassette (without frameshifts). Images were captured at 10 x magnification, g. Quantification of heterogeneous pan-neuronal splicing in adult day 3 RNAi-treated vs contr ol worms, **h.** Pan neuronal splicing index in hTau[P3OlL] vs WT control worms. Data are mean + s.e.m. of 3 biological replicates (n = 20-35 nematodes per group). Differences between conditions were assessed by two-way ANOVA while differences between genes or proteins were assessed by two-tailed t-tests (95% confidence interval). NS, no significance; *p<0.05, *\*\*p<* 0.01, ***p<0.001. The top 15 statistically significant up-regulated GO terms (related to Fig. lb) are shown in supplementary Fig,la. \A set of representative images of pan-neuronal splicing patterns (related to Fig. 1g) is included in supplementary Fig.lb-r.

ASEs allow a single eukaryotic gene to express functionally distinct proteins (*39, 40*) and belong to five distinct types: alternative 3′ splice site (A3SS), alternative 5′ splice site (A5SS), multiple exon skip (MES), intron retention (IR), and exon skip (ES). RNA-seq analysis of ASEs in the hTau.P301S transcriptome revealed that IR events were the most frequent ASE type (44 IR events in 106 total ASEs), ES events were slightly less frequent (42), while the number of the remaining three ASE types was much lower (A3SS, 7; A5SS, 7; MES, 6) (**Fig. 1c, Table S3**). Furthermore, GO analysis revealed distinct functional enrichments, such that A3SS-, MES-, and IR-events were enriched in genes involved in RNA metabolic processing (**Fig. 1d**), while A5SS events were enriched in genes that regulate mitochondrial ribosomes or relate to mitochondrial translation and respiratory system process (**Fig. 1d**). Some genes carrying A5SS and ES events co-located and regulated synaptic function (**Fig. 1d**). These results suggest that ASEs, especially A5SS and ES events, in genes related to mitochondrial and synaptic function were dysregulated in hTau.P301S mice.

As ES was a top event of ASE in hTau.P301S mice as well as its important role in neuronal function, we generated a roundworm *Caenorhabditis elegans* model to monitor neuronal ES event during ageing and in Tau pathology. The roundworm *Caenorhabditis elegans* strain KH2566, which carries a pair of pan-neuronally-expressed reporter minigenes, was used to examine whether and how aging impacts mRNA splicing and ASEs. In KH2566 worms, expression of the *C33H5.18* exon 2 fluorescent reporter minigenes is driven by the neuronal *rgef-1* promoter, and each of the minigenes carries a distinct frameshift mutation. ASEs in the reporter genes are monitored by fluorescence imaging to detect the intensity and ratio of GFP:mCherry fluorescence. Expression of GFP depends on the presence of exon 2, while expression of mCherry occurs when exon 2 is skipped (**Fig. 1e**). Therefore, the GFP:mCherry ratio is a quantitative measure of ASE events in neurons of KH2566 worms (**Fig. 1f**). The RNA splicing index (the ratio of green to red fluorescence in KH2566 worms) correlates inversely with the frequency of exon 2 skipping in worms carrying reporter minigenes. To test the system, worms were neuronal-specifically RNAi knocked down of a comprehensive set of conserved spliceosome components (**Fig. 1g; Table F 1b-r; Table S4**) (*41*). Reduction of these 13 conserved spliceosome components greatly increased exon 2 skipping, as evidenced by reduced GFP/RFP ratios (**Fig. 1g**); these data confirm that the KH2566 reporter gene system is an effective tool for monitoring the fidelity of endogenous mRNA splicing in the neurons of living *C. elegans*. This reporter gene system is also ideal for monitoring whether and how aging or AD-like disease impacts the fidelity of mRNA splicing. To use this reporter gene system in worms with tauopathy-like disease, KH2566 worms were crossed with CK12, hTau[P301L] worms (*42*) (*43*), a well-characterized strain expressing pan-neuronal human Tau 4R1N P301L that mimics tauopathy-like disease including impaired memory functions (*17, 44*).

Pre-mRNA splicing is tightly regulated during *C. elegans* development (*45, 46*) and studies using an mRNA splicing reporter gene suggest that some developmentally-regulated ASEs are expressed in a tissue-specific manner (*47*). In hTau[P301L] worms carrying the reporter minigenes described above, exon 2 skipping (ES) events increase consistently from adult day 1 (will be simplified to “day x” onwards) to day 3 and beyond with age, while in WT worms exon 2 skipping decreases from day 1 to day 3 before beginning to increase consistently with age (**Fig. 1h**). Notably, fewer exon 2 skipping events were observed in adult day 1 hTau[P301L] worms than in WT worms, whereas this relationship reversed, and more exon 2 skipping events were observed in hTau[P301L] worms than in WT worms after day 3 (**Fig. 1h**). We observed that the splicing index in WT nearly two-fold higher than in hTau[P301L] worms on day 3, but the difference decreased gradually from day 3 to day 13, when the differences were smaller (**Fig. 1h**). These results suggest that neuronal RNA splicing homeostasis is maintained in young hTau[P301L] worms by a compensatory mechanism. Interestingly, the splicing index showed different trends in the gut than in the central nervous system (CNS) of WT and hTau[P301L] worms, (**Table F 1s-u**), indicating that the compensatory mechanism in neurons is a tissue-specific phenomenon.

### NR upregulates genes involved in RNA splicing and components of the spliceosome in tauopathy

Previous studies show that mRNA processing is modulated by NAD^+^ in viruses, yeast, bacteria, and humans. Notably, mRNA splicing is perturbed in AD patients resulting in alterations to splicing signals, trans-acting regulatory factors, and components of the splicing machinery (*48–54*). Furthermore, a recent study suggests that aberrant RNA splicing processes contribute significantly to pathophysiology in the PS19 transgenic mouse model of tauopathy (*6*). Here, the effects of NR on RNA splicing were investigated in worm and mammalian cell-based tauopathy models.

WT and hTau[P301L] worms were treated with NR (1mM, 2mM, 5mM) or vehicle (control) from egg hatching on day 1 to day 15, were monitored for changes in fluorescence, and the splicing index was calculated for each experimental group. Interestingly, the splicing index increased more than 2-fold in NR-treated 1-day-old WT worms relative to vehicle-treated worms, but this increase was reduced by approximately 25% by day 2 (**Fig. 2a**). In contrast, 2mM NR only increased the RNA splicing index by 20% in 1-day-old hTau[P301L] worms, and the splicing index was not altered by NR in any other tested group of hTau[P301L] worms (**Fig. 2b**). These results suggest that NAD^+^ may only regulates RNA splicing during the earliest stage of worm development.

**Fig. 2.**
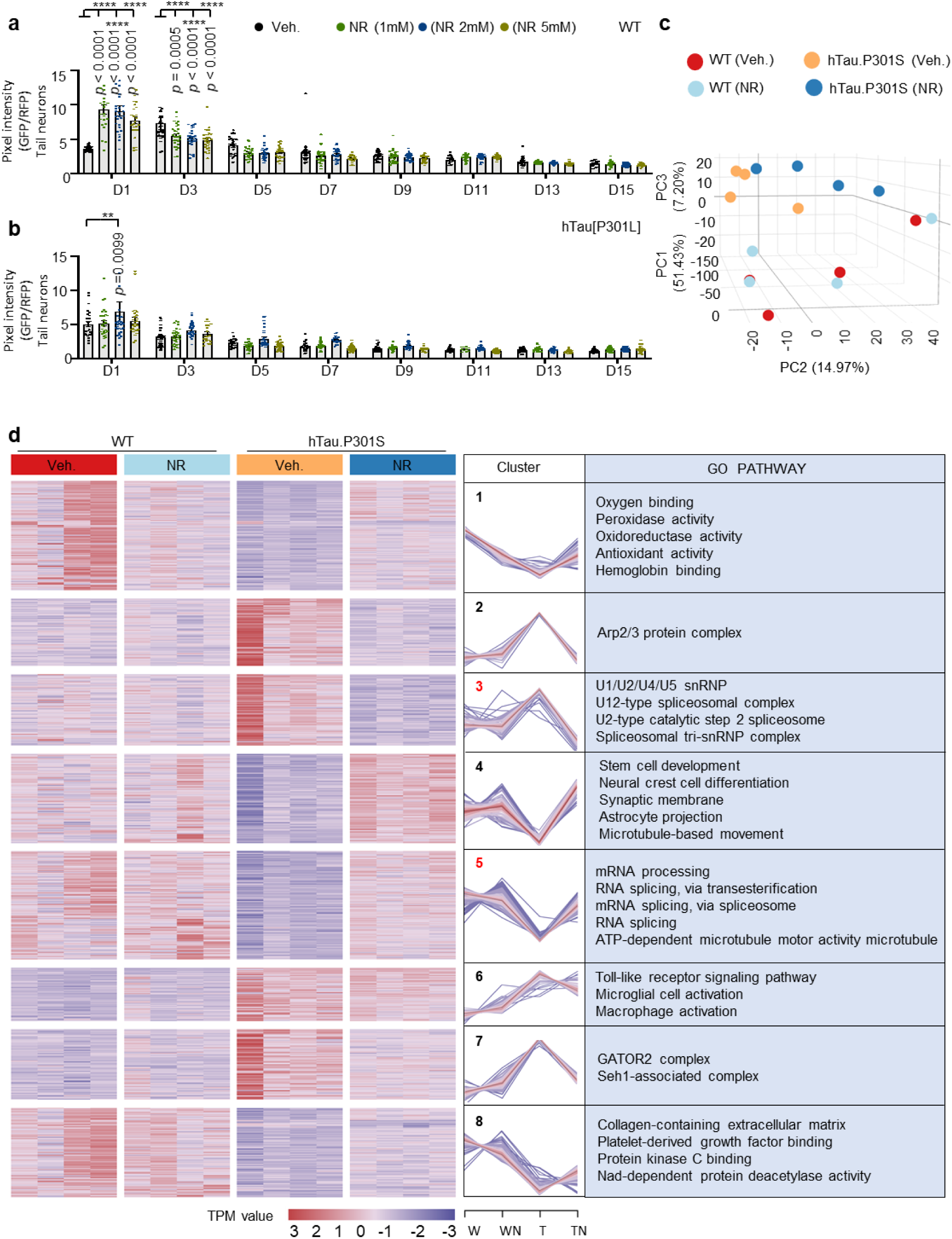
NR normalizes gene expression in tauopatbic worms. **a, b.** Pan neuronal RNA splicing index in day 1 to day 13 worms treated with 0, 1. 2, or 5 inM NR; **WT (a).** hTau[P3OlL] **(b).** Vehicle control data are for one set of experiments in Fig. Ih. c, 3D graph of PCA analysis of mouse hippocampal gene expression in WT (Veh.), WT (NR), hTau.P301S (Veh.), and hTau.P301S (NR) mice, d, Heatmaps show up– and down-DEGs in 4 experimental groups of mice as indicated (left panel). Eight clusters of DEGs (left panel) are represented graphically (middle panel). GO analysis of DEGs stratified by gene cluster (right panel). The criterion for statistical significance was adjusted P-value <0.05. GO terms shown in red are related to mRNA splicing or the spliceosome. Data are mean + s.e.in. of 3 biological replicates (n = 20-35 nematodes per group), one-way ANOVA was used to assess statistical significance. NS, no significance; *p<0.05, *\*\*p* < 0.01. *\*\*\*p* < 0.001.

RNA-seq data from the hippocampal region of mouse brains were used to determine whether and how NR influences mRNA transcription and splicing in hTau.P301S and WT mice. We treated both WT and hTau.P301S mice with NR (6 mM in drinking water, similar as we did before(*55*)) for 2 months followed by hippocampal tissue collection for RNA sequence. To identify general similarities in the RNA-seq datasets, principal component analysis (PCA) was conducted comparing hTau.P301S with or without NR to WT mice with or without NR. Results of PCA showed clear separation between transcriptional patterns in WT and hTau.P301S transgenic mice (**Fig. 2c; Table S5**). PCA also detected two subgroups of NR-treated hTau.P301S mice (**Fig. 2c**), but only partial separation between NR-treated and vehicle-treated WT mice (**Fig. 2c**). These data suggest that mRNA transcription and splicing in the brain hippocampal region may be significantly different in hTau.P301S and WT mice, possibly with hTau.P301S mice being more sensitive than WT mice to the effects of NR.

RNA-seq data were then used to identify genes and signaling pathways that were differentially-expressed in hTau.P301S or WT mice with or without NR. This analysis identified 730 up-or down-regulated differentially expressed genes (DEGs) (adjusted *p* values < 0.05) shown graphically with a heat map in **Fig. 2d (Fig 2d, left panel; Table S6**). These DEGs align into eight clusters which show different temporal expression patterns (**Fig. 2d, middle panel; Table S6**). Functional enrichment analysis revealed functions related to RNA-splicing in cluster 3 including 4 categories strongly related to the spliceosome, a large RNA-protein complex upon critical for pre-mRNA splicing. These categories were U1/U2/U4/U5 snRNP, U12-type spliceosome complex, U2-type catalytic step 2 spliceosome, and Spliceosomal tri-snRNP complex (*56*). Cluster 5 includes three functions directly related to pre-mRNA splicing and one related to ATP-dependent microtubule motor activity. Interestingly, the expression patterns were oppositely regulated in clusters 3 and 5, with79 genes in cluster 3 up-regulated in hTau.P301S mice relative to WT, and then rapidly down-regulated with NR treatment, while 120 genes in cluster 5 were down-regulated in hTau.P301S mice relative to WT but strongly up-regulated by NR. Notably, in WT mice, genes in clusters 3 and 5 were neither up-or down-regulated by NR (**Fig. 2d, right panel; Table S6**). These results suggest that genes directly involved in RNA splicing were downregulated in tauopathies. Furthermore, we propose that a compensatory mechanism induces the spliceosome in the absence of NR, but this is reversed in the presence of NR. As a result, compensatory up-regulation of spliceosome-related genes is absent in the hippocampal brain region of NR-treated hTau.P301S mice.

### Effect of NR on global transcriptome structure in tauopathic and WT mice

Figure 2 panels (c) and (d) show that Tau pathology polarizes the global transcriptome such that hTau.P301S and WT transcriptomic data map to opposite ends of the principal component 1 (PC1) and PC2 axes (Fig. 2c). NR remodels these relationships, which is consistent with the idea that NR modulates mRNA metabolism in tauopathic mice (Fig. 2d). This observation was explored further in the following experiments, which focus on quantitative expression of genes with open reading frames (see Methods for details) and the mechanisms underlying NR-dependent changes in patterns of gene expression. This analysis identified eight clusters (Fig. 3a**, Table S7**) with the most significant changes in expression profiles in transgenic hTau.P301S mice. These eight clusters were further grouped into three classes as follows: Class 1 clusters 1, 3, and 6 are characterized by upregulation of transcripts in NR-treated hTau.P301S mice relative to vehicle-treated hTau.P301S mice; Class 2 clusters 4 and 5 are characterized by downregulation of transcripts in NR-treated hTau.P301S mice relative to vehicle-treated hTau.P301S mice; Class 3 included only Cluster 8 and showed no transcriptomic changes between NR-treated and vehicle-treated hTau.P301S mice (Fig. 3b**, Table S8**).

**Fig. 3.**
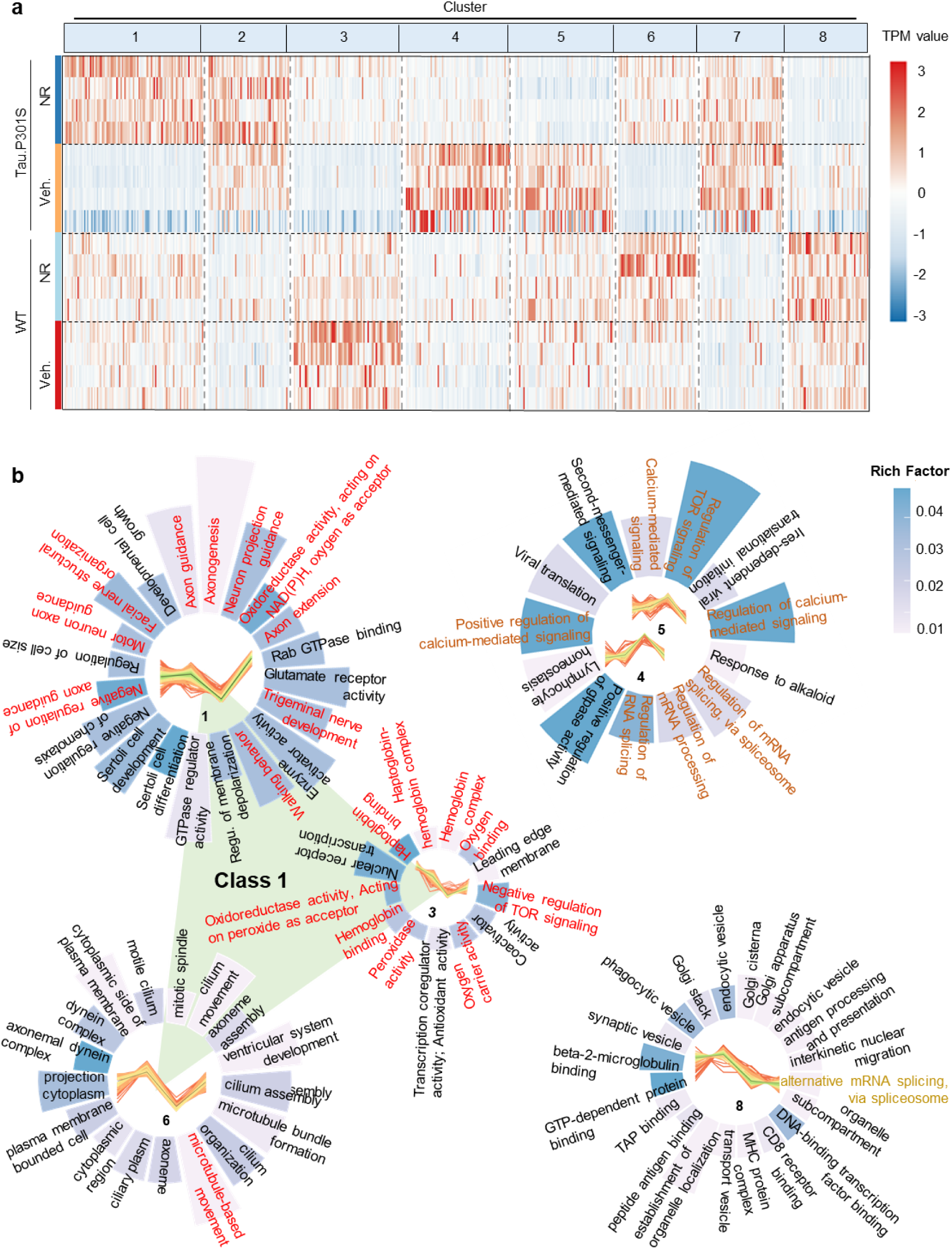
NR induces transcription of genes involved in axon development, oxygen metabolism, mitochondrion localization, and autophagy in tauopathic mic. **a.** Heatmaps showing DEGs and 8 classes of DEG derived by c-class cluster analysis for the 4 experimental groups of mice as indicated. The scale of relative differential expression is shown to the right, where red corresponds to maximum upregulation and blue corresponds to maximum downregulation of gene expression, b, GO terms enriched in each of the eight classes of clusters in **a;** each circular bargraph represents one cluster or class of cluster’s and bar length correlates with the number’ of DEGs represented. GO terms shown in red font relate to mRNA. Clusters 4 and 5 are similar and are grouped together. Class 1 includes clusters 1. 2, and 6 marked with red. Class 2 includes cluster’s 4 and 5 marked with orange. Class 3 includes cluster 8 marked with yellow.

Class 1 clusters, where NR stimulated expression of genes that are downregulated in hTau.P301S mice, were enriched in transcripts from 50 signaling pathways including 8 pathways related to nerve development and function. At the gene level, 75% of the genes played roles in axon genesis, extension, and guidance, suggesting possible involvement in formation of neural circuits and synapse cones. The latter processes could promote synthesis of Aβ, hyper-phosphorylation of tau, and AD pathogenesis (*57*). A pathway involved in trigeminal nerve development, a putative risk factor for dementia (*58*) is also represented Class 1 clusters. Eight pathways involved in oxygen metabolism were enriched in Class 1 clusters. Because functional oxygen metabolism is critical for cognitive function, the affected pathways could play roles in AD-associated synaptic dysfunction, neuronal death and tissue thinning in critical areas of the brain. Gene expression changes also affected microtubule-facilitated movement, mitochondrion localization, and transportation. Proper mitochondrial localization is required for formation of functional neural connections (*59*). In addition, the mammalian target of rapamycin (mTOR) pathway was downregulated by NR in Class 1 clusters, correlating with NR’s role in autophagy/mitophagy induction(*17, 60*); this could ameliorate AD pathology and symptoms (*61*). Expression of transcripts involved in facial nerve structure and organization was also altered by NR, which could potentially normalize facial and eye movement patterns in AD patients. Lastly, NR normalized expression of transcripts related to walking, which can be affected in the early stages of dementia (*62*); this is in line perfectly with our recent findings showing robust anti-ataxia capacity by NAD^+ (*55*)^, a finding further validated in a clinical trial(*63*). These results suggest that multiple mechanisms and pathways mediate the effects of NR on taupathic mice, including nerve development, oxygen metabolism, microtubule-facilitated movement, and autophagy (Fig. 3b**, Table S9**).

Class 2 clusters, where NR suppressed expression of genes that are upregulated in hTau.P301S mice, were enriched in 13 signaling pathways including three pathways related to calcium signaling. Excessive Ca^2+^ in mitochondria can lead to mitochondrial dysfunction, high levels of reactive oxygen species, increased apoptosis, and accelerated progression of AD pathology (*64*). Lastly, three pathways related to RNA splicing were enriched in Class 2 clusters (Fig. 3b**, Table S10**).

Class 3 clusters demonstrated no significant transcriptomic changes in NR-treated vs vehicle-treated mice. On the whole, these data suggest that NR normalizes deficiencies affecting the nervous system, oxygen metabolism, mitochondrial localization, autophagy, behavior, Ca^2+^ abundance and mTOR signaling in hTau.P301S mice. Thus, mRNA splicing is normalized in NR-treated hTau.P301S mice and the previously active compensatory mechanism becomes superfluous (Fig. 3b**, Table S11**).

### Overview of genes affected by ASEs and the effect of NR on ASE subtype and distribution in tauopathic mice

Alternative mRNA splicing is a critical source of splice-switching targets (*65, 66*) and is thought to play an important role in AD– and tauopathy-related disease progression. To understand this in greater detail, the number and subtype of ASEs were determined in hTau.P301S and WT mice with or without NR. A total of 267 ASEs were identified primarily belonging to MES, IR, or ES subtypes (Fig. 4a**-c**). Of these, 48 were identified by comparing NR-treated and vehicle-treated hTau.P301S mice and 21 were specific to NR-treated mice (Fig. 4a**, Table S12**). Fifty-three ASEs were identified by comparing NR-treated and vehicle-treated WT mice, and 37 were specific to NR-treated WT mice (Fig. 4a**, Table S12**). One hundred and six (106) ASEs were identified by comparing vehicle-treated hTau.P301S and vehicle-treated WT mice, and 73 of these were specific to hTau.P301S mice (Fig. 4a**, Table S12**). Interestingly, 20 ASEs were differentially expressed in vehicle-treated hTau.P301S vs. vehicle-treated WT mice and NR-treated hTau.P301S vs. vehicle-treated hTau.P301S mice, but not in NR-treated WT vs. vehicle-treated WT mice or NR-treated hTau.P301S vs. vehicle-treated WT mice. This indicates that treatment with NR selectively suppresses 20 ASEs that are specific to hTau.P301S mice (Fig. 4a**, Table S12**). One of these ASEs occurs in *Eva1-C*, whose pattern of expression suggests a role in axon guidance (*29*). This encouraged us to further explore the possible role of *Eva1-C* in tauopathic disease progression in the mouse.

**Fig. 4.**
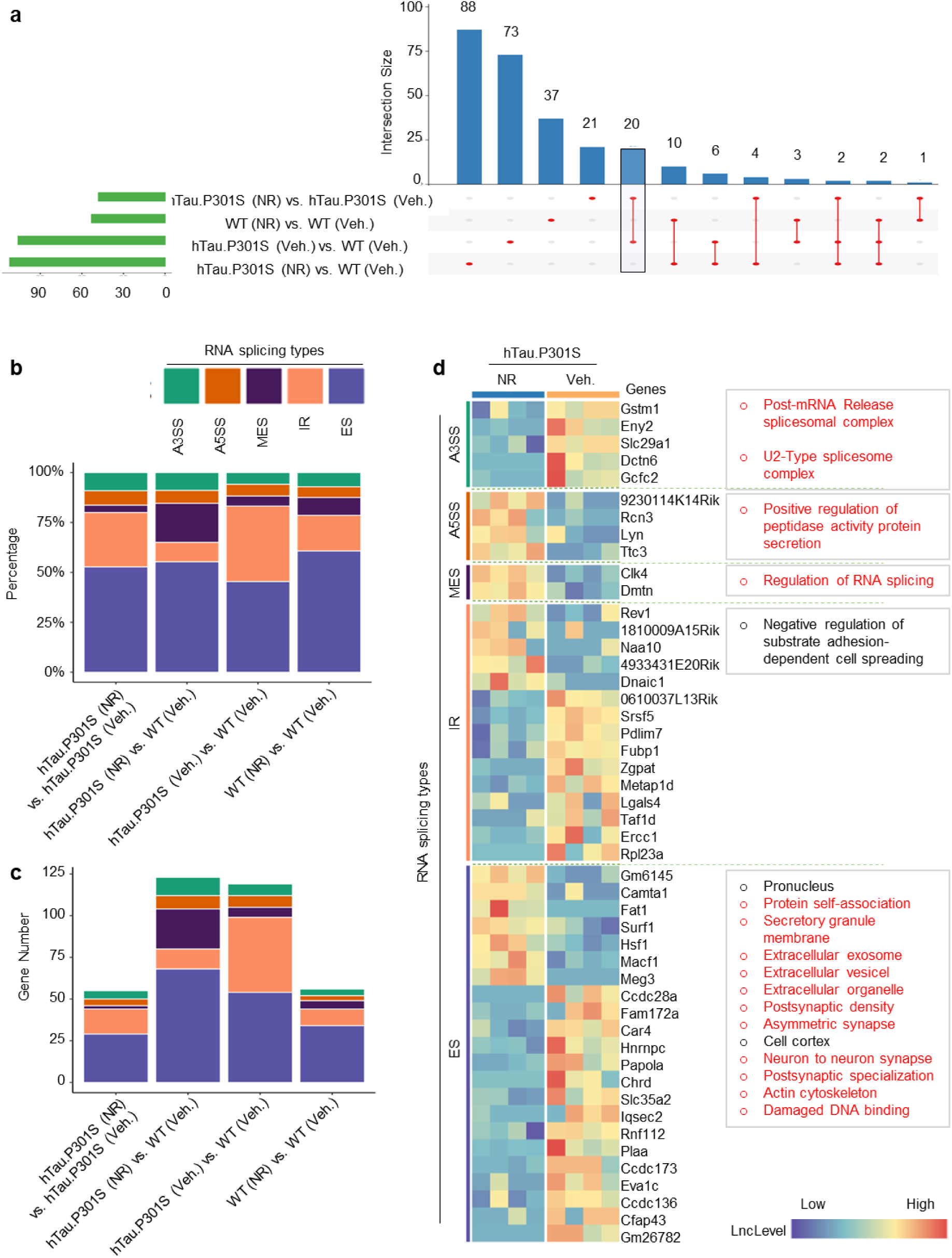
NR ameliorates multidimensional RNA splicing in Tauopathy. **a.** The UpSet plot in **(a)** summarizes the results of 6 pairwise and 2 three-way comparisons of ASEs affecting DEGs between hTau,P3OlS or WT mice treated with or without NR. The intersection size is the number of shared events between the experimental groups, b. For each comparison between experimental groups, the percent representation of each ASE subtype is represented by the height of the corresponding color-coded stacked bar. c. Sarne as (b) except bar height represents gene number instead of percentage of total events for each ASE subtype, d, Heatmap comparing DEGs in hTau,P3OlS transgenic mice in the presence vs. the absence of NR. where DEGs are further stratified by ASE subtype. Statistically significant enrichment of GO terms is shown for each DEG subtype. The criterion for statistical significance was adjusted P-value <0.05). Corresponding data for WT mice are shown in Extended Data Fig. 2.

We observed that A3SS and A5SS events represented a similar proportion of all ASEs in all experimental groups (Fig. 4b**; Table S13**). In addition, the majority of ASEs were detected in NR-treated hTau.P301S transgenic mice, indicating that targeting ASEs may be a beneficial effect of NR (Fig. 4c**; Table S13**).Thus, RNA-seq data were mined for ASEs specific to NR-treated hTau.P301S mice (Fig. 4d**; Table S14**). This analysis revealed five gene isoforms affected by A3SS events in NR-treated hTau.P301S mice including the post-mRNA release spliceosomal complex and the U2-Type spliceosome complex. The post-mRNA release spliceosomal complex catalyzes disassembly of the spliceosome separating U2, U5, U6, NTC (NineTeen Complex, CPX-1885), and the lariat-intron. The U2-Type spliceosome complex removes introns from pre-mRNA (*67*). Furthermore, A5SS events were detected in *Riken, Rcn3, Lyn*, and *Ttc3*, which regulate peptidase activity. This is consistent with earlier reports that peptidase activity is altered in AD (*68, 69*) and could implicate MES events and altered peptidase activity in AD progression. Figure 4b shows that the largest proportion of ASEs and the largest number of genes represented are ES events.

These results also show that ASEs in 22 genes affecting 54 isoforms were disproportionately represented in NR-treated vs vehicle-treated hTau.P302S mice and one of these genes is *Eva1-C*. To understand their significance, GO analysis was performed on this subset of genes. This analysis identified 13 core processes and pathways, 11 of which have potential involvement in AD pathophysiology. For example, our data show NR modulates protein self-association. Protein self-association leads to proteinaceous deposits, β-amyloid peptide-containing plaques, and neurofibrillary tangles by overcoming unfavorable electrostatic repulsions (*70, 71*). Furthermore, NR also modulates expression of genes related to extracellular organelles (*i.e.*, exosomes and vesicles); extracellular organelles carry disease-associated protein aggregates and are linked to AD pathology (*72*). The intracellular secretory granule, which has been reported to release CAP37, which is expressed at a higher level in the brains of patients with AD (*73, 74*); *CAP37* is a member of this subset of genes. Notably, GO terms related to “synapses” and “actin cytoskeleton” were also enriched in this subset of genes, which may relate to the fact that the actin cytoskeleton regulates the structure and function of dendritic spines (*75*). Genes that promoting the response to DNA damage are linked to AD progression (*76–78*); NR is also implicated in regulating ES events in these genes (Fig. 4d**; Table F 2**). These data suggest that NR facilitates multilayered control of ASE events in hTau.P301S mice and that additional research on the underlying mechanism is warranted.

### NR modulates splicing of Eva1-C pre-mRNA and improves cognitive function in tauopathic mice

To better understand the mechanism(s) by which NAD^+^ regulates RNA splicing, we focused on 22 transcripts whose expression and splicing are modulated by NR in hTau.P301S mice (Fig. 5a**, Table S15**). Because 15 of these transcripts encode proteins, the remaining 7 non-coding transcripts were excluded from further study. A comprehensive literature review identified five NR-responsive genes, *Fubp1*, *Zgpat*, *Zfp827*, *Tra2a*, and *C2cd5*, whose primary function appeared to involve binding to DNA, RNA, or other regulatory ions or biomolecules (*79*). Other genes in this group include *Adamts10, Zdbf2, Cacna1g*, and *Snrnp70*, whose expression patterns are altered in AD (*42, 80-83*). Furthermore, expression and splicing of *Eva1-C* transcripts are specifically regulated by NR in tauopathic mice (Fig. 4d) and EVA1-C is thought to play critical roles in neurons (*29*), although its involvement in AD pathology is not well understood; thus we focused on EVA1-C.

**Fig. 5.**
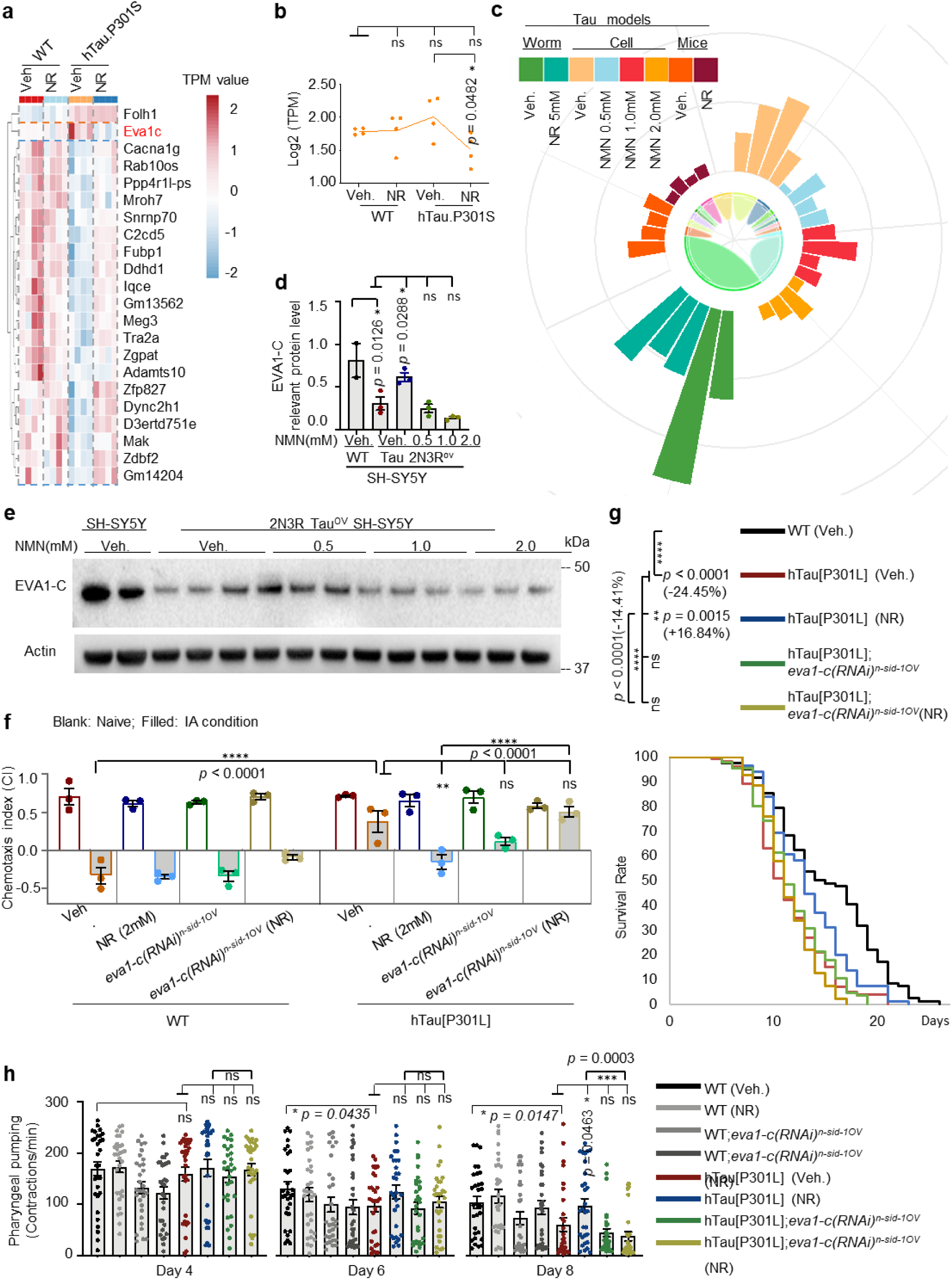
NR dependent expression of *eval-c* normalizes cognition and extends health span in tanopathic worms. **a.** Heatmap of ORF-containing DEGs affected by ASEs in hTau.P301S and WT mice treated with or without NR. **b.** Graph shows relative differential expression of *Eval-C* in the four indicated experimental groups, c, Circular bargraph representing expression *oĩEVAl-C* in worms (+/− NR), human cells (+/− NMN), and mice +/− NR), **d** f, Immunoblot analysis of EVA1-C protein in SH-SY5Y and 2N3R tau overexpressing SH-SY5Y cells treated with 0.5, 1.0 or 2.0 mM NMN or vehicle control. Quantification of **φ,e)** is shown in **(d.f). g, h**, Chemotaxis index was measured in WT, hTau[P3OlL], hTau[P3OlL] worms treated with NR. hTau[P3OlL] fed with *eval-c* RNAi, and hTau[P3OlL] fed with *eval-c* RNAi and NR. Lifespan was estimated from one set of experiments with three technical replicates (see Supplementary Table 17 for details) and 90-120 worms for each biological repeat. Data are shown as mean + s.e.m. Log-rank test was used to evaluate statistical significance (g). Pharyngeal pumping rate was measured in adult worms (n= 30 from 3 independent biological repeats) on days 4. 6. and 8 (h). Data from 3 biological replicates in e (n = 20-35 nematodes per group) are sho™ as mean + s.e.m. Two-way ANOVA was used to assess statistical signficance. NS, no significance; *p<0.05, **p<0.01. ***p< 0.001.

To clarify the roles and contribution of EVA1-C to AD pathology and progression, we used several tauopathic models including human SH-SY5Y cells overexpressing 2N3R Tau, hTau[P301L] *C. elegans*, hTau.P301S mice, and human brain tissue. The 2N3R-overexpressing SH-SY5Y cells were chosen because *in vitro* data indicate that phosphorylated recombinant 2N3R Tau is strongly associated with AD (*84*). We also used two *In vivo* AD models, hTau[P301L] *C. elegans* and hTau.P301S mice, as well as human brain samples in different Braak stages. The results show that *Eva1-C* mRNA is more abundant in vehicle-treated hTau.P301S mice than in vehicle-treated WT mice, but the abundance of *Eva1-C* mRNA decreased significantly in NR-treated hTau.P301S mice, while NR did not significantly alter expression of *Eva1-C* mRNA in WT mice (Fig. 5b). A similar decrease in abundance of *eva1-c* mRNA occurred in worm– and cell-based experimental models treated with NAD^+^ precursors nicotinamide mononucleotide (NMN) or NR (Fig. 5c**; Table S16**). In contrast, EVA1-C protein was approximately 2-fold higher in vehicle-treated control SH-SY5Y cells than in vehicle-treated 2N3R-overexpressing SH-SY5Y cells, and this difference was normalized when the cells were treated with 0.5mM NMN (Fig. 5d, e). However, no significant difference in EVA1-C protein expression was observed when the cells were treated with 1.0 or 2.0mM NMN (Fig. 5d, e). Because changes in Eva1-C mRNA and EVA1-C protein in these two experiments do not concur, the biotype of EVA1-C protein was examined (**Table F 3a, b; Table S17**). The results show that the upregulated *Eva1-C* transcript in hTau.P301S mice is a non-coding sequence, which could explain the discrepancy in experimental results in Fig. 5b, c vs Fig 5d, e.

The roles of EVA1-C in neurons and the effects of NR on expression of EVA1-C isoforms were investigated in hTau[P301L] worms with or without targeted RNAi knockdown of *eva1-c*. The results showed improved performance in memory tests in NR-treated vs vehicle-treated hTau[P301L] worms, but *eva1-c*-targeted RNAi abrogated this effect (Fig. 5f). Because previous studies suggest that AD patients have a shorter lifespan than healthy control groups (*85*), we also investigated the effects of NR and *eva1-c* knockdown on the lifespan of hTau[P301L] and WT control worms. Preliminary tests demonstrated that WT control worms had a longer lifespan than hTau[P301L] worms (Fig. 5g), which is in line with published results (*85, 86*). NR (2mM) increased the lifespan of hTau[P301L] worms by 16.84%, but did not extend the lifespan of *eva1-c*-knockdown hTau[P301L] worms or *eva1-c* knockdown WT control worms (Fig. 5g**; Table S18**), suggesting that *eva1-c* is a determinant of lifespan in these worm strains. We further measured pharyngeal pumping in NR-treated and vehicle-treated worms and established that treatment with NR did not change this parameter, a marker of healthspan (Fig. 5h). In summary, these data argue NR-induced lifespan extension and the improvement of memory-like capacity is dependent on EVA1-C.

### NR promotes expression of an EVA1-C isoform with high affinity for HSP70

We further designed experiments to explore the pathway(s) and effectors downstream of EVA1-C in the presence or absence of NR in tauopathic mice. Our approach was to mine genome-wide transcriptomic data for information on differential ASEs and combine the results with predictive modeling of protein 3D structures (using the AlphaFold2 model) and predicted PPIs.

First, ASEs and expression of alternative isoforms of *Eva1-C* were quantified in WT control and hTau.P301S mice in the presence or absence of NR (Fig. 6a**; Table F 3c**). The results reveal differential splicing of the middle exon of *Eva1-C*. Specifically, the middle exon of *Eva1-C* (90887491:90890939) is differentially included in WT and differentially excluded in hTau.P301S mice (Fig. 6a**, top panel; Table F 3c**). To confirm this result, RNA-seq data were analyzed using the mixture-of-isoforms (MISO) software. This analysis revealed that the IncLevel (probability of inclusion) of exon 2 was 0.74 vs 0.34 in WT vs hTau.P301S mice, respectively (Fig. 6a**, top panel; Table F 3c**). Furthermore, the right exon (90876190) was differentially excluded in NR-treated hTau.P301S mouse brain (IncLevel 1.0) and differentially included in vehicle-treated hTau.P301S mice (IncLevel 0.45) (Fig. 6a**, bottom panel; Table F 3c**). The relative abundance of all *Eva1-C* isoforms (n=6) in all experimental groups is summarized in Fig. 6b. Two isoforms are non-protein-coding transcripts (denoted as “NR*” to distinguish with the NAD^+^ precursor “NR”) and four are protein-coding transcripts (NM) (Fig. 6b). Interestingly, the noncoding transcripts were significantly downregulated by NR, suggesting differential expression of active EVA1-C in the presence of NR. Interestingly, the protein-coding 2362 bp transcript was not detected in mouse brain (Fig. 6b). Furthermore, expression of the 2366 bp transcript was higher in NR-treated WT and hTau.P301S mice than in the respective vehicle-treated control mice (Fig. 6b); in contrast, NR decreased expression of the 2045 bp and 2222 bp transcripts in hTau.P301S mice (Fig. 6b). These results could have implications regarding relative expression of functional EVA1-C protein in NR-or vehicle-treated tauopathic or WT mice.

**Fig. 6.**
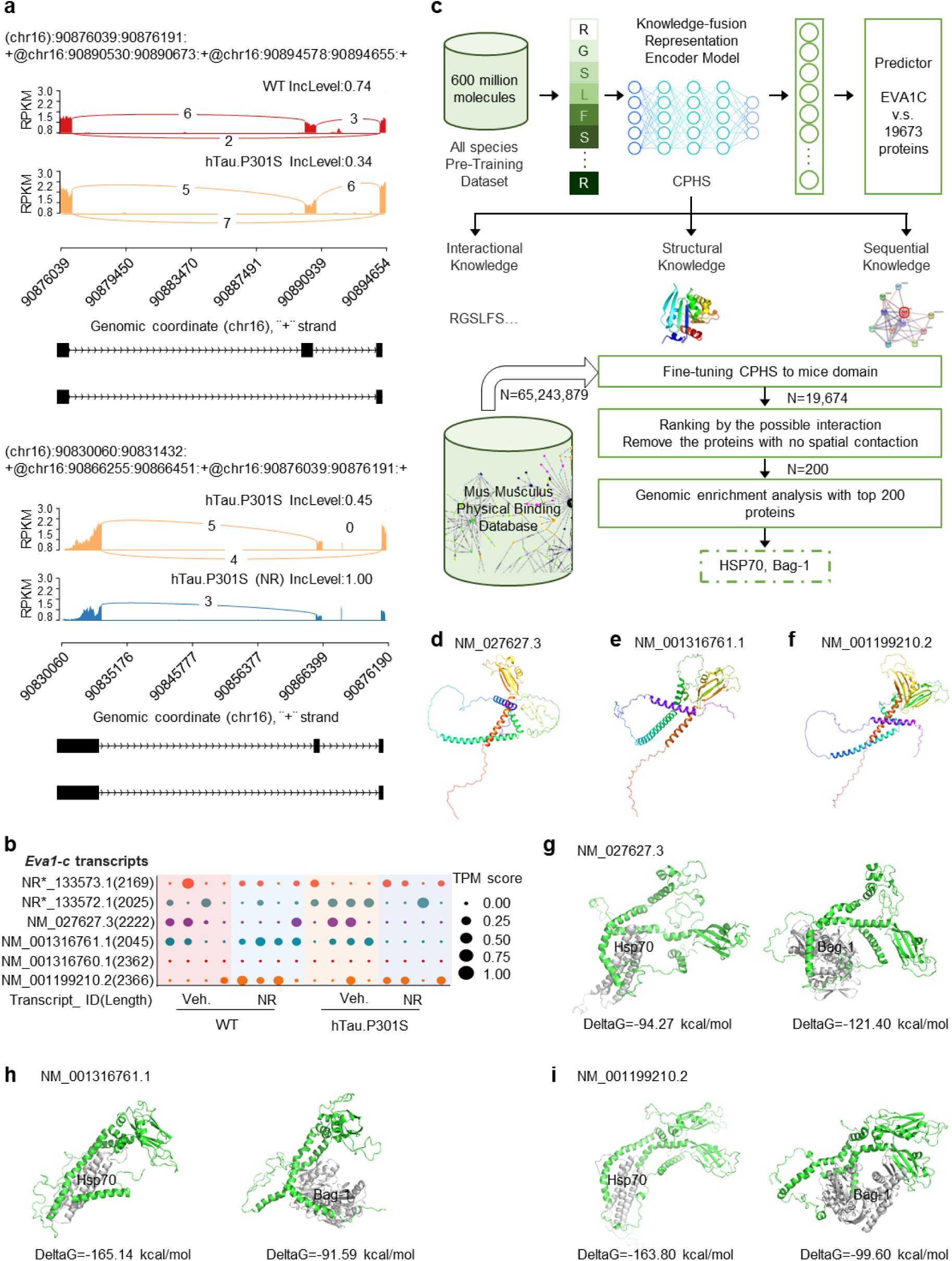
Deep learning strategies to characterize EVAI C protein protein interactions in hTau.P3OIS and WT mice. **a.** Sashimi plot of ASEs in *Eval-C* in WT, hTau.P301S, and hTau.P301S+NR mice. Diagrams show the read coverage of exons (upper) with incLevel shown above corresponding diagram. AS models are shown schematically below (lower panel). The curve indicates the splicing pattern and the number above the curve is the number of corresponding RNA-seq reads. Similar analysis of an individual sample is shown in Extended Data Fig. 3c. RNA-Seq data from WT, hTau.P3OIS, and hTauP3OlS + NR mice are represented in red. orange, and blue, respectively. Reads Per Kilobase Million (RPKM) are plotted against reads split across the splice junction (junction depth) to appear as arcs representing the number of splice junction connected exons per sample. Alternative isoforms are shown below’ the junction tracks, **b.** Dot plots representing the relative abundance of 6 *Eval-C* transcript isoforms in four experimental conditions, c. Schematic outline to illustrate processes of model pre-training and virtual screening. Upper panel: the pre-training dataset was analysed using the Knowledge-fusion Representation Encoder Model, and interactional, structural, and sequential knowledge was used to train the EVA1-C bonding predictor within the top 0.003%. Bottom panel: virtual screening was performed using the Mus Musculus Physical Binding database containing 65.243,879 molecules. The pretrained model was fine-tuned using mouse domains with 19674 sequences. The top 100 putative protein-interactors were identified and ranked while proteins with no structural contact were removed from consider ation. Genomic enrichment analysis was used to identify two lead hits. HSP70 and BAG-1, **d-f**, Predicted 3D structures of three EVA1-C isoforms generated with AlphaFold2. **g-i.** Protein docking simulations for EVA1-C (green) and HSP70 (gray, purple, blue) or BAG-l(gray, purple, blue). Additional details including IDDT and pTMscore are shown in Extended Data Fig. 4. The list of top 100 pr oteins is shown in Supplementary Table 19.

Next, we used a novel deep-learning AI-based approach to predict the EVA1-C PPI network and potential downstream effectors of EVA1-C in tauopathic models. This was accomplished primarily using the AlphaFold2 software to predict EVA1-C isoform-specific 3D structures from the EVA1-C primary amino acid sequence (*87*) (Fig. 6c**-f**). The top structural models were then identified and used in protein docking simulations to identify putative PPIs of interest.

This analysis focused on three EVA1-C isoforms: NM_001316761.1, NM_001199210.2 and NM_027627.3. The top five protein conformations for each isoform were identified after the conformations were ranked based on the Local Distance Difference Test (IDDT) (*88*) and the template modeling score (TM-score) (*89*). Each of the protein conformations with highest IDDT value and TM-score for each isoform were selected for further analysis. For the three selected protein conformations, >80% of the structure had IDDT >0.6, 40% of the structure had IDDT >0.75, and all three structures had TM-scores >0.5 (**Table F 4**). Potential PPIs were evaluated for all three isoforms, and proteins that showed high potential PPIs with all three isoforms were selected for further study. Potential PPIs were fine-tuned using the STRING-mouse species dataset (*90*) as well as Deep PPI (*34*), GNN-PPI (*91*), PIPR (*92*) and ONTOPROTEIN (*93*) for comparison. Using the STRING-mouse species dataset, F1 scores were 79.26 ± 0.2 and 87.82 ± 0.4 for the Breadth First Search (BFS) and Depth First Search (DFS) split training and evaluation datasets, respectively, where higher F1 score indicates better performance. Our model outperformed Deep PPI, GNN-PPI and PIPR (**Table 1**).

**Table 1:**
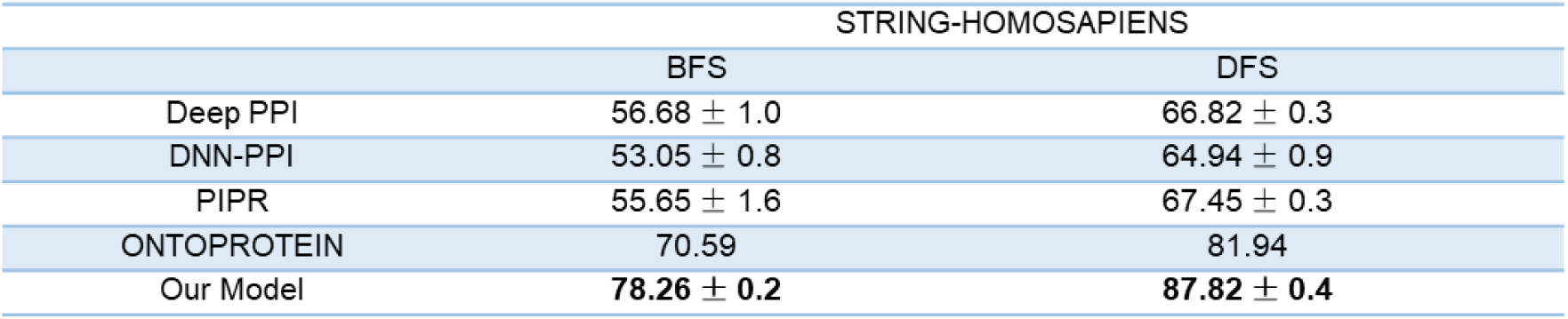
A comparison of performance between our model and other PPI Prediction Models on STRING-HOMOSAPIENS.

|  | STRING-HOMOSAPIENS |  |
| --- | --- | --- |
|  | BFS | DFS |
| Deep PPI | $56.68 \pm 1.0$ | $66.82 \pm 0.3$ |
| DNN-PPI | $53.05 \pm 0.8$ | $64.94 \pm 0.9$ |
| PIPR | $55.65 \pm 1.6$ | $67.45 \pm 0.3$ |
| ONTOPROTEIN | 70.59 | 81.94 |
| Our Model | **$78.26 \pm 0.2$** | **$87.82 \pm 0.4$** |

After fine-tuning the model with the mouse species dataset, PPIs were predicted for the three EVA-1C isoforms. After the Ankyrin repeat, SOCS box protein (ASB) and the leucine-rich repeat-containing (LRRC) protein were removed due to likely biological irrelevance, HSP70 ranked first in three predictions and BAG-1 ranked in the top 100 (**Table S19**). Because HSP70 is a critically important molecular chaperone and it interacts with BAG-1 to induce autophagy (*94, 95*), subsequent studies focused on interactions between EVA1-C, HSP70 and BAG-1 (Fig. 6c).

Protein docking simulations were performed to estimate and compare binding modes and energetics using predicted models of the three isoforms of EVA1-C docked with BAG-1 or HSP70. The top 10 of 3000 energy-minimized conformations were identified (Fig. 6g**-i**). The predicted binding affinities (ΔG) of HSP70 to NM_001316761.1, NM_001199210.2 and NM_027627.3 isoforms of EVA1-C were –163.80 kcal/mol,-165.14 kcal/mol, and –94.27 kcal/mol, respectively (*p* < 0.05). For BAG-1, ΔG values for binding NM_027627.3, NM_001316761.1 and NM_001199210.2 were –121.40 kcal/mol, –99.60 kcal/mol, and –91.59 kcal/mol, respectively (*p* < 0.05).Notably, in hTau.P301S mice, NR promotes differential expression of the NM_001199210.2 isoform of EVA1-C, which has higher affinity for HSP70 than for BAG-1 (ΔG= –165.14 vs. –91.59).

### NR/NMN induces *Bag-1* mRNA and regulates the interaction between BAG-1 and HSP70

We then investigated Eva1-C-dependent effects on transcription of *Hsp70* and *Bag-1* in NR-or vehicle-treated tauopathic and WT worms. We crossed CK12 (hTau[P301L]) worms with TU3401 worms (characterized with hypersensitive neuronal RNAi by feeding), followed by feeding the crossed worms with *eva1-c* RNAi to generate a transient hTau[P301L];*eva1-c(RNAi)^n-sid-1OV^* worm line. The TU3401 worms carry mutated *sid-1* in all tissues except neurons, where functional SID-1 was overexpressed, confering neuron-specific hypersensitivity to exogenous RNAi (*96*). Notably, transcripts of *C. elegans hsp6*, a homolog of *Hsp70* and *Bag-1* increased approximately 2-fold in NR-treated transgenic hTau[P301L] worms. While RNAi-mediated knockdown of *Eva1-C* did not interfere with NR-dependent stimulation of *Hsp6* transcription, knockdown of *Eva1-C* reduced the NR-dependent increase in *Bag-1* transcription by approximately 50% (Fig. 7a). Thus, NR modulates transcription of *Bag-1* in an *Eva1-C*-dependent manner, while NR modulates transcription of *Hsp6* in an *Eva1-C*-independent manner.

**Fig. 7.**
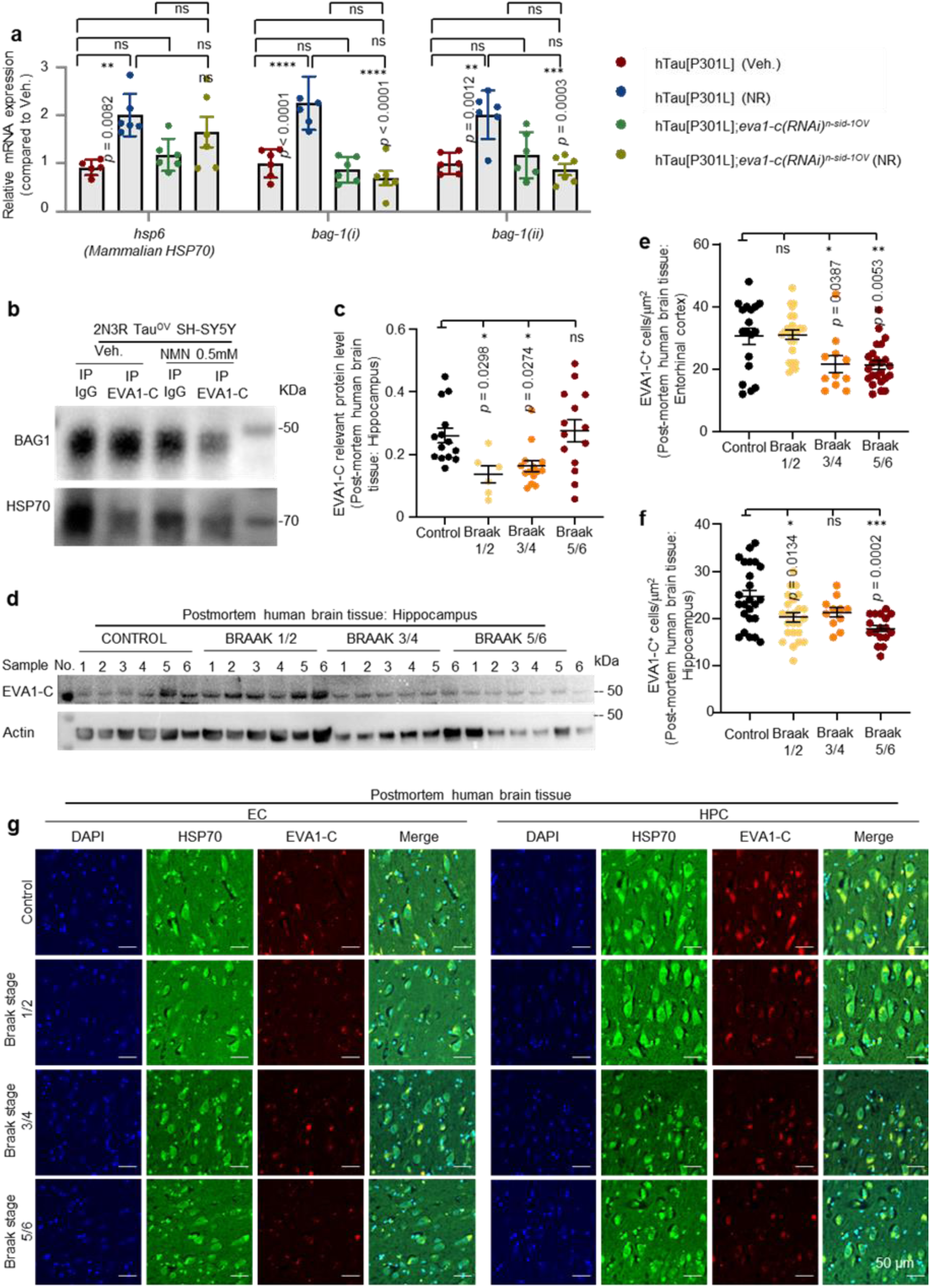
NR and disease progression modulate expression of EVA1-C and binding to BAG-1 and HSP7O. **a.** Graphs show the relative expression of *hsp6* and *bag-1* in day 1 adult IrTau[P3OlL] and WT worms incubated with or without evαf-c-targeted RNAi and with or without NMN. The data shown in **a** are from one of two biological replicates from a single representative experiment (1 biological and 3 technical repeats) horn a total of 2 biological replicates, b. Co­immunoprecipitation (Co-IP) assays investigate the interactions between EVA1-C and HSP7O or BAG-1, c,d. Western blot shows expression of EVA1-C in hippocampal brain from AD patients with different Braak stages and age-matched healthy control (6-14 samples per group), and quantification of (b) is shown in (c). e-g, The number of EVA1-C^-^ cells from AD patients at different Braak stages of disease and age-matched healthy controls (10 samples per group). Representative samples from errtorhirral cortex (e) and hippocampal (f) brain regions were analyzed. Representative images of immuno fluorescent signals detecting EVA1-C (red). HSP7O (green), and DAPI (blue) in control, Braak stage 1/2, Braak stage 3/4, and Braak stage 5/6 (gh). Data are shown as mean + s.e.m.; ns. no statistical significance; *p<0.05, **p<0.01, *\*\*\*p<*0.001.Statistical significance was assessed by two-way or one-way ANOVA followed by Šidák’s multiple comparisons test as appropriate. All experiments were per formed at least twice.

Using SH-SY5Y cells overexpressing 2N3R Tau, the interactions between EVA1-C, HSP70 and BAG-1 were examined using Western blotting and immunoprecipitation assays. NR (0.5 mM) slightly downregulated expression of BAG-1, while it did not alter expression of HSP70 in 2N3R Tau-overexpressing SH-SY5Y cells (Fig. 7b). Co-immunoprecipitation (co-IP) assays also indicate an NR has a more potent impact on the interaction between EVA1-C and BAG-1 than on the interaction between EVA1-C and HSP70 (Fig. 7b**; Table F 5a**). Our data suggest HSP70 is the most susceptible protein in PPI with EVA1-C in tauopathy. Thus, to better understand the interaction between EVA1-C and HSP70 in human cells, we explored whether EVA1-C and HSP70 are co-expressed and co-localize in human brain hippocampal tissue from AD patients and normal controls.

For this purpose, total lysates were prepared from hippocampal regions of post-mortem human brains from patients assigned to different Braak stages or normal control patients. Immunoblotting data show that EVA1-C is less abundant in brain tissue from patients at Braak stages 1/2 or 3/4 than in normal controls; data in Brakk stages5/6 were with high variation (Fig. 7c, d**; Table F 5b**). As here the immunoblotting data were reflections of the average EVA1-C protein levels in all brain cell types, we asked changes of EVA1-C levels specifically in neurons across Braak stages in hippocampus, as well as in the entorhinal cortex (the first region of Tau pathology in AD, see review (*97*)). We observed around 30% reduction in number of EVA1-C immunoreactive (EVA1-C^+^) neurons in the entorhinal cortex of patients at Braak stages 3/4 and 5/6 relative to cognitive normal controls (Fig. 7e, g). Similarly, the number of EVA1-C^+^ neurons in hippocampal brain tissue was significantly lower in patients at Braak stages 1/2 and worsen at 5/6 cells compared to controls (Fig. 7f, g). We observed co-localization between EVA1-C and HSP70 in neurons (Fig. 7g). Detailed information of AD patients and their cognitive normal controls are provided (**Table S20**). Collectively, these data suggest that NR increased transcriptional levels of *hsp-6* and *bag-1* and in an EVA1-C-dependent manner; the human brain neuronal EVA1-C was reduced crossing Braak stages in both the entorhinal cortex and hippocampus.

## Discussion

This study demonstrates that NR-stimulated differential expression of *Eva1-C* isoforms and events downstream of EVA1-C contribute to the neurological benefits of NR in experimental models of AD-like tauopathies. The study employs a multipronged approach involving genome-wide transcriptomic data mining, 3D protein structure modeling and computational protein docking and PPI network prediction, and the computational output was tested in mouse, worm and human cell-based models for tauopathy. The results suggest that differential regulation of ASEs contributes significantly to the therapeutic efficacy of NR and NMN for treating AD. The results of this study also argue in favor of future development of splice-switching therapeutics for AD and AD-like diseases.

We reported that NR-regulated ASEs have potential as therapeutic targets for AD, we also identified genes and ASE-dependent isoforms of these genes that are regulated by NR in tauopathic model systems. In this study, transcriptomic analysis of hippocampal mouse brain tissue and a pan-neuronal splicing reporter worm were used to demonstrate dysregulation of ASEs in tauopathic worms. This result is consistent with results of a recent study on PS19 mice (*6*). We also demonstrate that neuronal ASEs are dysregulated with increasing age of WT worms in a manner distinct from dysregulation of ASEs associated with pan-neuronal expression of hTau 4R1N. A previous study reported similar results in the gut of aged worms (*41*).

To understand the potential of NAD^+^ to manipulate ASEs for therapeutic purposes, transcriptomic data were mined for information on the biological functions and GO pathways susceptible to regulation by NAD^+^-stimulated ASEs. The results identified pathways associated with axon genesis, axon extension, axon guidance, oxygen metabolism, mitochondrion localization, and autophagy, all of which are primarily induced by NAD^+^ in models of tauopathy. Several previous studies reported that different isoforms of the same functional genes have distinct or opposite functions in the neuronal system, including mGlu3R, RNCAM, TRIP8b, QKI, Grm1, RARbeta, EAAT2, Vegfa, BDNF, plectin (*98–105*). Furthermore, the ratio of some isoforms of these genes is altered by NAD^+^ in tauopathic animals, as observed in the present study. We speculate that ASE-regulated genes functionally related to axon development play a critical role in tauopathy and that these genes represent promising targets for NAD^+^-based splice-switching therapeutics for AD.

We found that ASEs in genes involved in RNA splicing are disproportionately dysregulated in tauopathies. This leads us to speculate that the beneficial effects of NR in tauopathies may be mediated by its ability to normalize ASEs in these genes. We also found disproportionate NR-dependent effects on A3SS and MES subtypes of ASEs and disproportionate ES events affecting neurons, phenomena that may be important areas of future study. Thus, the prevalence of different ASE subtypes should be considered when designing precision splice-switching therapeutic strategies.

Our transcriptomic data mining identified *EVA1-C* splicing as a potentially important target of NR in tauopathies. Previous studies reported that EVA1-C was necessary for heparin-binding (*106*) which could relate to charge compensation during aggregation of Tau protein (*107*). In addition, a prior study in mice indicated that expression of EVA1-C played a role in axon guidance in the mouse nervous system (*29*). Here, we discovered that EVA1-C was dramatically reduced in AD patients, in a paten crossing Braak stages in both the entorhinal cortex and hippocampal neurons. We also show that NR stimulated expression of EVA1-C in an isoform-specific manner and increased EVA1-C protein in the brains of hTau.P301S mice.

Computational approaches were used to predict critical PPIs downstream of EVA-1C in taupathic models. This analysis led us to focus on interactions between EVA1-C, HSP70 and BAG-1, subsequently validated in selected experimental models of tauopathy. BAG family proteins are co-chaperones of HSP70. BAG1 can negatively regulate protein refolding by HSP70 *in vitro* and *in vivo* (*94, 108-110*). It was proposed that BAG1/BAG3 switching acts as a link between macroautophagy and the ubiquitin-proteasome system (*95, 111, 112*). However, little is known about whether and how the BAG1/BAG3 balance influences AD pathophysiology. We speculate NR may regulate the BAG1/BAG3 balance in an EVA1-C isoform-dependent manner in tauopathy; further experiments are necessary. Our further mechanistic studies show that NR/NMN reduced ES events in *Eva1-C*; NAD^+^ supplementation increased expression of an EVA1-C isoform with higher affinity for HSP70. Notably, NR modulated expression of *Bag-1* mRNA in an *Eva1-C*-dependent manner.

The present study is subject to some limitations. For example, the available bulk RNAseq database is only representative of gene expression values that are averaged over different conditions. Second, EVA1-C isoform-specific antibodies were not available for use in the present study, which in turn ruled out immunologically based approaches to verify isoform-specific outcomes. To compensate, AI-based deep learning approaches were used to simulate EVA1-C isoform-specific interactions and outcomes. Therefore, further investigation is warranted to verify EVA1-C isoform-specific interactions with HSP70 and BAG1. Despite these limitations, the present study provides novel and important insights into NAD^+^-dependent regulation of ASEs in tauopathy and suggests important areas for future development of ASE-targeted therapeutics for AD– and AD-like diseases. At the same time, NAD^+^-based therapeutics continue to be relevant for modulating other pathways involved in AD pathogenesis including proteostasis, energy metabolism, immune response, glial function, and neuronal death (*16, 17, 113*). Intriguingly, several NAD^+^-related clinical trials targeting AD are in progress (NCT05617508 and NCT04430517), highlighting the importance of further in-depth mechanistic studies. This study contributes to improved understanding of mechanisms underlying transcriptional diversity, it could point the way towards successful future development of ASE-targeted precision medicine for treating AD and related dementias.

## Materials and Methods

### Bioinformatics and RNA seq analysis

#### Quality control for raw FASTQ data

Standard next-gen DNA sequencing protocols were used to derive DNA primary sequences from raw data. FASTQ format files were generated containing sequence reads and sequencing quality information. Software fastqc and trim_galore were used to remove adapters and N-containing and low-quality reads (<Q20). RNA-seq data from 16 mice were processed and analyzed.

#### Acquisition and alignment of gene-level expression data

Mouse reference genome mm10 and the GTF format gene annotation file were downloaded from the UCSC website (https://hgdownload-test.gi.ucsc.edu/goldenPath/mm10/bigZips/). Then STAR was used to index the reference genome mm10 and align the clean FASTQ files. Bam files were generated from aligned data using the Samtools software and bam files were analyzed using RSEM. RSEM was used to calculate and map number of reads and raw counts to corresponding genes. Then, raw counts were corrected for gene length and influences of sequencing depth and gene length on read count were accounted for. Last, we calculated Fragments Per Kilobase of the exon model per Million mapped fragments (FPKM).

#### Analysis of differential expression and gene enrichment at gene-level

Pairwise difference comparisons among WT, WT+NR, hTau.P301S, and hTau.P301S +NR (n=4 per group) were performed with R package DESeq2 (v1.16.1) (*114*). Significance criteria for differentially expressed genes were P adjust <0.05 and abs (log2 (fold change))>0.5. Average FPKM values per group were analyzed, gene expression trends determined, and the fuzzy c-means method was used to cluster with normalized values (*115*). GO enrichment analysis and KEGG enrichment analysis were performed using R package clusterProfiler (4.0)(*114*). Significance criteria for enrichment of GO and KEGG terms were p adjust<0.05 (i.e., Benjamini & Hochberg correction).

#### Transcript acquisition and analysis at transcript-level

STAR was used to align data from different groups (*116*) and RSEM was used for quantitation (*117*). Count and TPM at the transcript level were analyzed with Deseq2. ClusterProfiler and Mfuzz were used to analyze GO and KEGG term enrichment as described previously.

#### Acquisition and analysis of alternative splicing events

FASTQ files were analyzed using RSEM and all five subclasses of ASEs were analyzed using rMATs. Absolute values of lncLever greater than 0.1 and Padjust < 0.05 were considered statistically significant. GO and KEGG enrichment for ASEs per group was also analyzed.

#### AI-based prediction and modeling of protein structures and PPIs

An AI-based deep-learning method and pre-training were used to leverage cross-species multi-level structural data and obtain a knowledge-fusion model. The model was refined using mouse data and interaction fields to obtain task-desired representations.

Protein Prompt Learning: In-context learning leverages extra semantic tokens to steer the model towards task-specific representations. To extend this concept to proteins, we employed pre-training task-specific tokens called prompts into the protein sequence, whereby each dedicated sentinel token is associated with a pre-training task and can be leveraged to coax the model towards task-specific protein representations. A prompt-tuning module was used during model refinement. This allowed us to combine prompts on-demand and bridge the gap between pre-trained knowledge and downstream tasks.

The hierarchical-structure-aware model, PromptProtein, was used as a backbone. PromptProtein utilizes three modules, objectives-masked language modeling, alpha-carbon coordinate prediction, and physical-binding prediction, to acquire primary, secondary, tertiary, and quaternary structural knowledge and the corresponding prompts, including protein sequence [SEQ], protein 3D structure [CRD], and protein protein interaction [PPI]. To evaluate this model, we used the Protein-Protein Interaction (PPI) benchmark generated by Multifaceted protein–protein interaction prediction based on Siamese residual RCNN | Bioinformatics. The two methods, Breath-First Search (BFS) and Depth-First Search (DFS), were used to split training and evaluation datasets. The F1 scores were used as a criterion to compare performance PromptProtein with Deep PPI (*34*), GNN-PPI (*91*), PIPR (*92*) and ONTOPROTEIN (*93*) and demonstrate the superiority of our model. Note that the original model is trained on datasets that are not species-specific. We first pre-finetuned parameters of the whole model for mouse using the AlphaFold Protein Structure Database, which contains 79,301 predicted structures and STRING which contains 22,048 sequences and 3,126,208 interaction pairs. Initially, the prompt-tuning module was only applied to the STRING dataset. After the initial stage of tuning, the model was used to predict PPI probabilities within the STRING dataset for mouse using EVA1-C as the query protein. Nuclear proteins and other proteins intrinsically unlikely to interact with EVA1C were removed, and the top 100 remaining potential interactors were evaluated manually (Fig. 6c).

#### Protein structure prediction, protein docking and molecular dynamics

Predicted EVA1-C protein structures were generated with ColabFold, which uses an accelerated protein structure prediction algorithm incorporating the fast homology search capability of MMseqs2 with the knowledgebase associated with AlphaFold2 and/or RoseTTAFold. Then, structures with the highest average pLDDT and pTMscore values were used for protein docking simulations and analyses of molecular dynamics.

Protein docking simulations used the ZDOCK server (https://zdock.umassmed.edu/) restricted to structures derived from X-Ray diffraction data resolved to <3 Å. In this study, EVA1C was docked with BAG-1 (PDB id: 3CQX) and HSP-70 (PDB id: 1i6z). Molecular dynamics simulations were performed for the top ten docked structures and average binding affinities (ΔG) were calculated.

Molecular dynamics simulations used the pdb4amber program to prepare and download original PDB files from the RCSB. The tags of all non-canonical residues were sorted and those with the fewest occurrences were merged into the protein structure. The complex files were inspected manually and the potential of the selected complexes as protein-ligand models was evaluated.

Using an RTX3080 GPU, simulations were run over approximately 30 hours allowing 100 ns per protein-ligand complex with the periodic boundary condition in the NPT ensemble. The detailed steps are: 1) 5,000 cycles of minimization (MAXCYC=5,000) in a solvated system using the steepest descent algorithm for cycles 1-2,500 and the conjugate gradient algorithm for the remaining cycles (MAXCYC – NCYC); 2) The NVT simulation was heated gradually from 0 to 303.15 K over 500 ps; 3) The heated system was equilibrated with the NPT ensemble over 1 ns; 4) Production simulation was run at 303.15 K at ambient atmospheric pressure and the SHAKE algorithm was used to constrain all covalent bonds involving hydrogen and does not calculate the forces of hydrogen bonds; 5) Finally, a 100 ns production simulation was performed collecting snapshots at 1 ps intervals. MM-PBSA was used to calculate 100 snapshots in the last nanosecond. Conformations with the highest binding affinities are represented graphically in Fig. 6**, g-i**. The most probable binding modes were deduced using 100ns molecular dynamics simulations and MM-PBSA.

### Human brain samples

#### Ethical

The study was performed in accordance with the Helsinki Declaration and Principles for Ethical Research. All participants gave informed consent in writing. The regional Ethics Committee for Medical Research in the South-East of Norway (REK 82685) and the local Data Protection Officer approved the study.

#### Western blot

Western blot assays were done as previously describe(*55*). Briefly, human brain samples were collected and firstly homogenized in tissue homogenizer (NextAdvance) with 0.5 mm zirconium oxide beads and ice-cold Triton X-100 buffer containing 1x protease inhibitor (catalog no. B14002; Bimake) and 1x phosphatase inhibitor (catalog no. B15002; Bimake). For homogenization, shark the tube at 4 °C at maximum speed for 5 min, and then centrifugation (21,000 × g) at 4°C for 10 min, the collected supernatant as soluble fraction. The pellet was washed 3 times with 1% Triton X-100 buffer, and solubilize the insoluble pellet with 1x radioimmunoprocipitation (RIPA) buffer containing 8 M Urea, 5% SDS, 1x protease inhibitor, and phosphatase inhibitor. Centrifugation (21,000 x g,) at 4°C for 10 min, and the supernatant was collected as insoluble fraction. The proteins concentration was tested via BSA method, and the insoluble fraction of protein sample (30 μg) was prepared and denatured at 95°C for 10 min and used in this study. NuPAGE 4-12% Bis-Tris Protein Gel (catalog no. NP0336BOX; Thermo Fisher Scientific) was used to separate proteins run at 200V 45 min. Transfer system was set at 25V for 30 min on PVDF membrane (for all the proteins from 15 kDa to 350 kDa).Various antibodies were probed. 5% BSA which dissolved in Tris-HCl buffer containing 0.1% Tween-20 (1x TBST) was using to block non-specific binding. Blots were incubation with Eva-1 Homolog C antibody (1:500) (Novus, catalog no. NBP1-88937), and actin (1:5000) (Invitrogen, MA1-744) at 4°C overnight, respectively. Blots were washed 3 times with 1x TBST,and incubated with either anti-rabbit IgG HRP-linked antibody (Cell signaling, 7074) or anti-mouse IgG HRP-linked antibody (Cell signaling, 7076) with dilution 1: 5000 at room temperature for 1h.. ChemiDoc XRS System (Bio-Rad Laboratories) was used to detect Chemiluminescence. Quantification was performed using ImageJ

#### Immunofluorescence/immunohistochemistry

Human brain sections were prepared from the prefrontal cortex, entorhinal cortex and hippocampus of patients designated Braak stage 1/2, Braak stage ¾ or Braak stage 5/6 and normal healthy controls. Only one brain region/Braak stage was sample per individual patient/control subject. Tissue sections (7 uM) were slide-mounted and then sequentially dehydrated in xylene, 50% ethanol, and H_2_O_2_. Subsequently, sections were rinsed 3 x 10 min in PB (125nM phosphate buffer, pH 7.4) three times. After pre-incubation in 5% BSA TBS-TX (50 mM Tris, 0.87% sodium chloride, 0.5% Triton X-100) for 1.5 h, the sections were incubated with primary antibodies (1:300 in TBS-TX) at 4 °C overnight. After rinsing, sections were incubated with secondary antibody (1:1000) and washed 3 x 10 min in PB. Slides were covered with the Prolonged Gold antifade with DAPI reagent (P36931, Invitrogen) and imaged using an automated slide scanner/fluorescent imaging system (AxioScan Z1, Zeiss). Digital images were captured with a 20X objective (NA 0.8). ZEN lite Blue software was used for data analysis and image quantification used a 5 μm 2 x 2 square grid equidistantly distributed over the brain region of interest. In each section and region, eight sampling areas were selected randomly and quantified. The EVA1-C/BAG-1 and EVA1-C/HSP70 co-localization index was quantified in a similar manner.

#### Cell culture for mechanistic studies

Human neuroblastoma SH-SY5Y and 2N3R Tau-overexpressing SH-SY5Y cells (generated by M. Akbari) were used in this study. The cells were maintained in GIBCO DMEM supplemented with 10% FBS with 1% penicillin-streptomycin (P&S) at 37°C under 5% CO_2_. After treatment, cells were collected for Western blot or Co-immunoprecipitation (CoIP) assay.

The CoIP assay was performed as previously described(*55*). Subsequent protein analysis via Western blot was carried out as described above using Eva-1 Homolog C Antibody (catalog no. NBP1-88937) (1:1,000 dilution, primary). Secondary antibody was diluted 1:5,000 in blocking buffer.

#### Co-immunoprecipitation (CoIP) assays

SH-SY5Y cells were trypsinized, washed twice with ice-cold PBS, released and collected by centrifugation. Cell pellets were flash frozen in liquid N_2_. and then resuspended in 2 PCV hypotonic lysis buffer (20 mM Hepes pH 7.9, 2 mM MgCl, 0.2 mM EGTA, 10% glycerol, 0.1 mM PMSF, 2 mM DTT, 1x Halt™ Protease Inhibitor Cocktail), placed in liquid nitrogen for 5 min, and then subjected to three freeze/thaw cycles. After the last freeze/thaw cycle, NaCl and NP-40 were added to final concentrations of 0.5 M and 0.5% (v/v) respectively, and then incubated for 20 min on ice. The samples were diluted with 8 PCV hypotonic lysis buffer containing 50 mM NaCl and sonicated. Nucleic acids were degraded by adding either 1.2 µl CaCl_2_ (2.5 M) and 25 U DNase or 2.1 µg RNaseI and 50 U MNase. Nuclease reactions were incubated for 1 h at 4 °C. Samples were centrifuged, supernatants collected, and aliquots of the supernatant (0.5 mg) were incubated with 2 µg anti-IgG antibody, anti-ELP1 (Abcam ab56362) or IgG (rabbit IgG, Diagenode, C15410206) or mouse IgG (Diagenode, C15400001) at 4 °C overnight. for purified FLAG-ELP1, AAG or Δ80 AAG, 1 µg protein was mixed with 1 µg antibody and incubated overnight at 4 °C in 200 µl final volume IP buffer (20 mM HEPES pH 7.9, 2 mM MgCl2, 0.2 mM EGTA, 10% (v/v) glycerol, 0.1 mM PMSF, 2 mM DTT, 140 mM NaCl, 0.01% (v/v) Nonidet-P40). Protein-A Dynabeads or Protein-A sepharose beads were equilibrated in IP buffer at 4 °C overnight, washed three times, added to samples and incubated for 4 h at 4 °C. The supernatant was removed and beads washed three times with cold wash buffer. Beads were boiled in Laemmli buffer and analyzed by Western blot as described.

#### C. elegans

**Table 2.** *C. elegans* strains and genetics.

| Strain genotypes | Source |
| --- | --- |
| KH2566: *ybEx2566 [rgef-1::C33H5.18E1E2(+1)E3-P2A-mCherry + rgef-1::C33H5.18E1E2(+1)E3-(+2)P2A-EGFP + pBSKII(-)]* | Kuroyanagi Hidehito |
| EFF084: *sid-1(pk3321) V;uls69 [pCFJ90 (myo-2p::mCherry) + unc-119p::sid-1] V;ybEx2566 [rgef-1::C33H5.18E1E2(+1)E3-P2A-mCherry + rgef-1::C33H5.18E1E2(+1)E3-(+2)P2A-EGFP + pBSKII(-)]* | The Fang Lab |
| *Is[Paex-3::tau4R1N(P301L), Pmyo-2::gfp]; ybEx2566 [rgef-1::C33H5.18E1E2(+1)E3-P2A-mCherry + rgef-1::C33H5.18E1E2(+1)E3-(+2)P2A-EGFP + pBSKII(-)]* | The Fang Lab |
| KH2235: *lin-15 (n765) ybIs2167 [eef-1A.1::RET-1E4E5(+1)E6-GGS6-mCherry eef-1A.1::RET-1E4E5(+1)E6-(+2)GGS6-EGFP lin-15 (+) pRG5271Neo] X* | Kuroyanagi Hidehito |
| EFF023: *Is[Paex-3::tau4R1N(P301L), Pmyo-2::gfp];ybEx2261 [Peef-1A.1::C33H5.18E1E2(+1)E3-GGS6-mCherry, Peef-1A.1::C33H5.18E1E2(+1)E3-(+2)GGS6-EGFP pBSKII(-)]* | The Fang Lab |
| TU3401: *sid-1(pk3321) V;uls69 [pCFJ90 (myo-2p::mCherry) + unc-119p::sid-1] V* | The Rafal Lab |
| EFF029: *Is[paex-3::tau4R1N(P301L) + pmyo-2::gfp];sid-1(pk3321) V;uls69 [pCFJ90 (myo-2p::mCherry) + unc-119p::sid-1] V* | The Fang Lab |

#### *C. elegans* drug treatment and RNAi experiments

*C. elegans* were cultured at 20°C on standard nematode growth medium (NGM) agar plates with OP50 unless stated otherwise. Nicotinamide riboside (NR) was custom synthesized by Maintain Biotech Company (1094-61-7) and prepared at 1, 2 or 5 mM in water. Worms were treated with drugs from day 1 (day of hatching) and. RNAi was delivered to worms on treatment plates.

#### Short-term memory assay

Chemotaxis towards an attractant (*i.e.*, isoamyl alcohol (IA) on a 10 cm agar plate was performed at 20 °C as previously described(*55*). More specifically, 200-300 synchronized day 1 worms were collected and washed 5 times with M9 buffer and placed on 6 cm NGM plates (without OP50) with IA for 120 min conditioning or without IA for 120 min. For pre-conditioning, 10 μl pure IA was placed in the middle of the plate lid at the start of the incubation period. Assay plates were prepared 30 min before starting the assay by adding 20 µl 20 mM NaN3, covering the ‘IA’-treatment area with a 0.5×0.5cm piece of Parafilm. Then, worms were collected in M9 buffer, quickly dried with tissue paper, placed on the ‘source point’, and 4 μl diluted IA (1/50) was added to the Parafilm. The testing plates were then quickly sealed with Parafilm. After 120 min, the number of worms in the vicinity of ‘source points’ S, IA, and ‘T’ was counted. The chemotaxis index was (#‘IA’ – #‘T’)/(#’IA’ + # ‘T’ + #‘S’), where ‘#’ denotes numbers (*118*). A lower test score correlates with higher memory capacity/performance. Three to five biological replicates were performed.

### Lifespan and pharyngeal pumping rate assays

#### Lifespan assay

Lifespan assays were conducted at 20°C on 3.5 cm NGM plates as described previously(*55*). Drugs were diluted in water (*i.e.*, 2 mM NR) before animals were dosed. Synchronous animals were generated via bleaching under defined conditions. Thirty-five to forty DL4 larvae per technical repetition were transferred to 3.5 cm plates. In one biological repeat. At least 100 synchronous worms from 3 technical repeats were tested. The plates were changed twice a week to ensure continuous exposure to the same drug dosage. Worms were scored daily and the number of living and dead worms was reccorded. Criteria for death were absence of pharyngeal pumping and unresponsive to touch. Worms that died due to gonad extrusion, internal bagging, or crawling on the edge of the plates were censored and given a weight of ‘0’, so they did not contribute to mortality calculations. The log-rank test (Mantel–Cox) was used to calculate the mean, the standard deviation of the mean, and the p-value. Kaplan Meier (K-M) analysis was used to derive survival curves. Statistical analyses were carried out using Prism (GraphPad Software).

#### Pharyngeal pumping rate

Animals were synchronized and maintained as described for the lifespan assay. The number of pharyngeal contractions in 30 s per adult worm was counted manually(*55*).

#### Quantification of *C. elegans* tissue mRNA

Real-time PCR was performed as previously described(*17*). Worms were collected and washed with M9 buffer and RNA was prepared using TRIzol™ (Thermo Fisher, Catalog number: 15596026) according to the manufacturer’s recommendations. cDNA was synthesized using iScriptTM cDNA Synthesis Kit (catalog no. 1708890, BioRad). Reverse transcription was carried out for 5 min at 25°C, 20 min at 46°C, 1 min at 95°C, and finally at 4 °C for storage. cDNA and transcript counts were determined using standard qRT-PCR protocols. All rections were run in triplicate. The primers used were:

*hsp-6*: 5’-TCGTGAACGTTTCAGCCAGA-3’ and 5’-CTCAGCGGCATTCTTTTCGG-3’

*bag-1*(I): 5’-ACCAAAGCAGGTCGAGATGG-3’ and 5’-CGTCTTGCGTTTTTCACGGT-3’

*bag-1*(II): 5’-CGCCACCGACAATGATGTTG-3’ and 5’-ATGAGCATTTTGAAGCCCGC-3’

*pmp-3*: 5’-ATGATAAATCAGCGTCCCGAC-3’ and 5’-TTGCAACGAGAGCAACTGAAC-3’

We used the QuantStudioTM 7 Flex System v1.1 (applied biosystems by Life Technologies). Cycle parameters were: 95 °C for 10 min, 40 cycles of 95 °C,15s; 60 °C,1 min. Melting curves were calculated after 95 °C for 15s, followed by 60°C for 1 min.

#### Transgenic mice

All animal procedures and protocols were approved by the Jinan University Institutional Animal Care and Use Committee. Animals were maintained and fed under standard conditions. The hTau.P301S mouse strain was a generous gift from Michel Goedert (Cambridge, United Kingdom). Behavioral tests were performed using 12-month-old mice. The mouse strain was constructed as described previously(*119*). The regional Ethics Committee for Medical Research in South-East Norway (FOTS ID 16060) and the local Data Protection Officer approved the study.

#### Hippocampal sample acquisition and mRNA extraction and detection

Hippocampus tissue from Thy1-hTau.P301S mice was resuspended in Trizol and dispersed by shaking. Aliquots were separated by agarose gel electrophoresis to ascertain RNA integrity and purity. NanoPhotometer spectrophotometry and the Agilent 2100 bioanalyzer were also used to confirm purity and integrity of RNA samples, respectively.

#### Library construction and quality control

Extracted mRNA (total ≥800ng) was used to build the library with Illumina’s NEBNext® UltraTM RNA Library Prep Kit. The Oligo (dT) magnetic beads were used to enrich for poly(A)-containing mRNA as input for library construction. To ensure the quality of the library, the effective concentration (no less than 2nM) was accurately quantified. DNA sequencing was performed according to Illumina DNA sequencing protocols.

#### Statistical analysis

All “wet-lab” data are presented as mean ± S.E.M., unless otherwise specified. A two-tailed unpaired t-test was used for pairwise comparisons between groups. Group differences were analyzed using one-way analysis of variance (ANOVA) followed by Šidák’s multiple comparisons test or two-way ANOVA followed by Tukey’s or Dunnett’s multiple comparisons test for multiple groups and determined to be normal distributions. For other datasets, Mann-Whitney U or Kruskal–Wallis tests were used. All statistical analyses used GraphPad Prism 8.0 software. The criterion for statistical significance was P value <0.05.

## Acknowledgements

E.F.F. is supported by Cure Alzheimer’s Fund (#282952)‘HELSE SØR-ØST (#2020001, #2021021, #2023093), the Research Council of Norway (#262175, #334361), Molecule AG/VITADAO (#282942), NordForsk Foundation (#119986), the National Natural Science Foundation of China (#81971327), Akershus University Hospital (#269901, #261973, #262960), the Civitan Norges Forskningsfond for Alzheimers sykdom (#281931), the Czech Republic-Norway KAPPA programme (with Martin Vyhnálek, #TO01000215), and the Rosa sløyfe/Norwegian Cancer Society & Norwegian Breast Cancer Society (#207819). R.X.A. and S.Q.Z were funded by the China Scholarship Council (http//:www.csc.edu.cn/); the funders had no role in study design, data collection and analysis, decision to publish, or preparation of the manuscript. Y.H. is supported by The Natural Science Foundation of China (No.82101664), and Medical Scientific Research Foundation of Guangdong Province of China (No.A2022007). Support for some of the transgenic nematode models used came from a Collaborative Innovation Award from the Howard Hughes Medical Institute to G.A.C. G.Y. is supported by the ERC IMI (101005122), the H2020 (952172), the MRC (MC/PC/21013), the Royal Society (IEC/NSFC/211235), the NVIDIA Academic Hardware Grant Program, and the UKRI Future Leaders Fellowship (MR/V023799/1). E.F.F. has an MTA with LMITO Therapeutics Inc (South Korea), a CRADA arrangement with ChromaDex (USA), and a commercialization agreement with Molecule AG/VITADAO, and is a consultant to Aladdin Healthcare Technologies (UK and Germany), the Vancouver Dementia Prevention Centre (Canada), Intellectual Labs (Norway), MindRank AI (China), and NYo3 (China). All other authors have nothing to declare. All data needed to evaluate the conclusions in the paper are present in the paper and/or the Supplementary Materials.

