## Supplemental Figure 1-5, Supplemental Table 4, 18,20. for "NAD^+^ - and EVA1-C-dependent reversal of neurological deficits is mediated by differential alternative RNA splicing in tauopathic animal models"

**Fig. S1.**

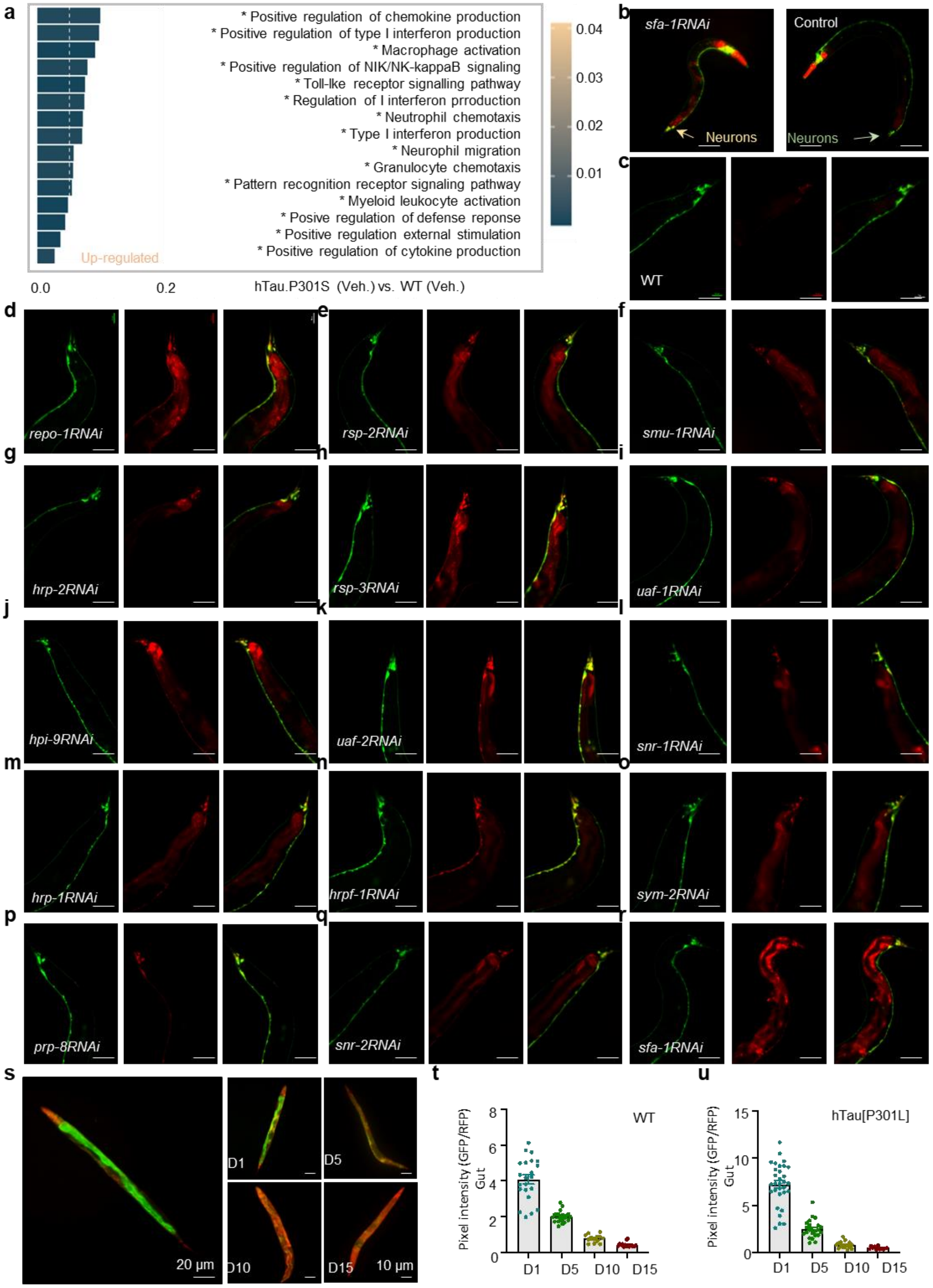
a
* Positive regulation of chemokine production
* Positive regulation of type I interferon production
* Macrophage activation
* Positive regulation of NIK/NK-kappaB signaling
* Toll-lke receptor signalling pathway
* Regulation of I interferon prroduction
* Neutrophil chemotaxis
* Type I interferon production
* Neurophil migration
* Granulocyte chemotaxis
* Pattern recognition receptor signaling pathway
* Myeloid leukocyte activation
* Posive regulation of defense reponse
* Positive regulation external stimulation
* Positive regulation of cytokine production
Up-regulated
0.04
0.03
0.02
0.01
0.0
0.2
hTau.P301S (Veh.) vs. WT (Veh.)
b
sfa-1RNAi
Control
Neurons
Neurons
c
WT
d
repo-1RNAi
e
rsp-2RNAi
f
smu-1RNAi
g
hrp-2RNAi
h
rsp-3RNAi
i
uaf-1RNAi
j
hpi-9RNAi
k
uaf-2RNAi
l
snr-1RNAi
m
hrp-1RNAi
n
hrpf-1RNAi
o
sym-2RNAi
p
prp-8RNAi
q
snr-2RNAi
r
sfa-1RNAi
s
20 µm
D1
D5
D10
D15
10 µm
t
WT
Pixel intensity (GFP/RFP) Gut
8
6
4
2
0
D1
D5
D10
D15
u
hTau[P301L]
Pixel intensity (GFP/RFP) Gut
15
10
5
0
D1
D5
D10
D15

**The role of mRNA splicing in Tau pathology and aging**

**a**, Top 15 most statistically significant upregulated Gene Ontology (GO) term pathways enriched in the gene list of upregulated mRNA transcripts (Fisher's adjusted P-values < 0.05) in mice. **b**, Neuron-specific rgef-1 splicing in day 1 *C.elegans* feeding with (left) and without (right) *sfa-1* RNAi. **c-r**, Heterogeneous splicing patterns in response to knockdown of conserved splicing factors in *C. elegans*. Control (feeding with L4440 RNAi) (**c**) and pan-neuronal knocked down of *repo-1* (**d**), *rsp-2* (**e**), *smu-1* (**f**), *hrp-2* (**g**), *rsp-3* (**h**), *uaf-1* (**i**), *hpi-9* (**j**), *uaf-2* (**k**), *snr-1* (**l**), *hrp-1* (**m**), *hrpf-1* (**n**), *sym-2* (**o**), *prp-8* (**p**), *snr-2* (**q**), and *sfa-1* (**r**) with RNAi from egg hatching, with images taken on day 3 of adulthood. **s**, A representative set of images of changes in RNA splicing in the gut tissue over the ages. **t, u,** Changes of splicing index (GFP/RFP) in gut between WT (t) and hTau[P301L] (u) worms of days 1, 5, 10, and 15.

**Fig. S2.**

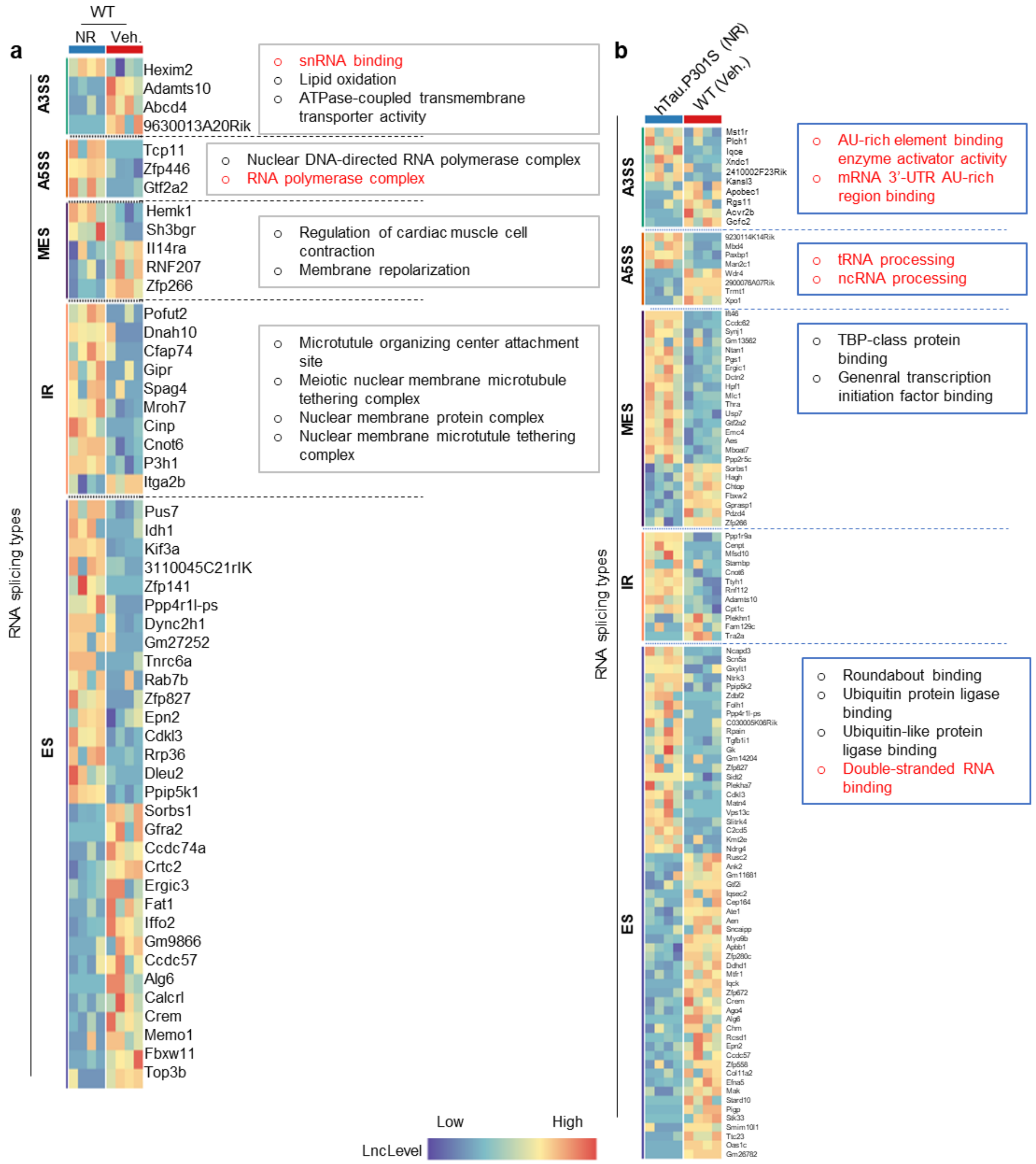


**Alternative RNA splicing events in different conditions**

**a-b**, Heatmaps comparing AS events based on alternative 3' splice site (A3SS), alternative 5' splice site (A5SS), multiple exon skip (MES), intron retention (IR), and exon skip (ES). Statistically significant Gene Ontology (GO) term pathways enriched in the gene list of changes in mRNA transcripts (Fisher's adjusted P-values < 0.05) in different types of alternative RNA splicing in mice. Heatmaps comparing AS events between WT mice treated with and without NR (**a**); data of hTau.P301S transgenic mice (NR) compared to WT (Veh.) were shown in (**b)**. Additional information was shown in Table S21, 22.

**Fig. S3.**

a

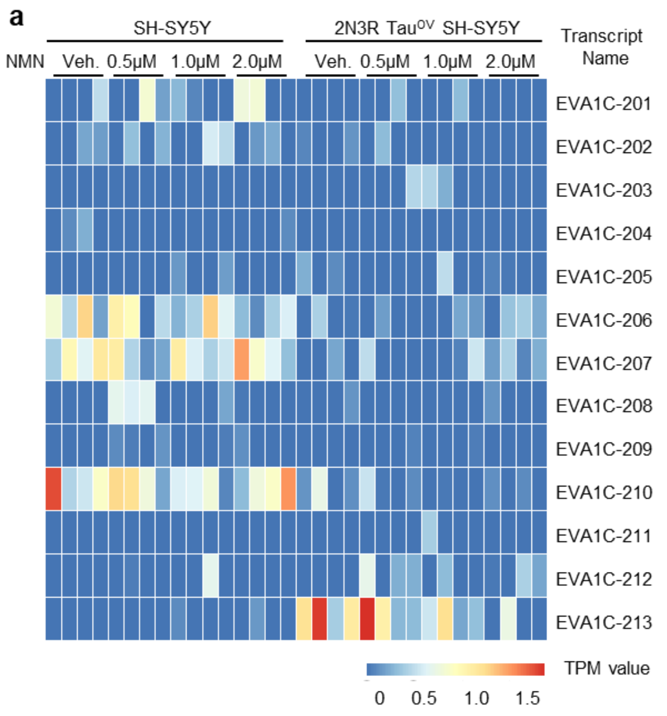


b

| Name | ID | Length (bp) | Protein | Biotype |
|---|---|---|---|---|
| *EVA1C -201* | ENST0000 0300255.7 | 1671 | 441aa | Protein coding |
| *EVA1C -202* | ENST0000 0382699.7 | 1668 | 438aa | Protein coding |
| *EVA1C -203* | ENST0000 0401402.7 | 1558 | 393aa | Protein coding |
| *EVA1C -204* | ENST0000 0412833.1 | 936 | 153aa | Protein coding |
| *EVA1C -205* | ENST0000 0435323.5 | 1601 | 70aa | Nonsense mediated decay |
| *EVA1C -206* | ENST0000 0437338.5 | 1745 | 70aa | Nonsense mediated decay |
| *EVA1C -207* | ENST0000 0457807.5 | 1548 | 192aa | Nonsense mediated decay |
| *EVA1C -208* | ENST0000 0459833.5 | 687 | No protein | Protein coding CDS not defined |
| *EVA1C -209* | ENST0000 0464037.5 | 2492 | No protein | Retained intron |
| *EVA1C -210* | ENST0000 0469079.5 | 1689 | No protein | Retained intron |
| *EVA1C -211* | ENST0000 0481638.1 | 384 | No protein | Protein coding CDS not defined |
| *EVA1C -212* | ENST0000 0485488.1 | 720 | No protein | Protein coding CDS not defined |
| *EVA1C -213* | ENST0000 0496615.5 | 492 | No potein | Protein coding CDS not defined |

c

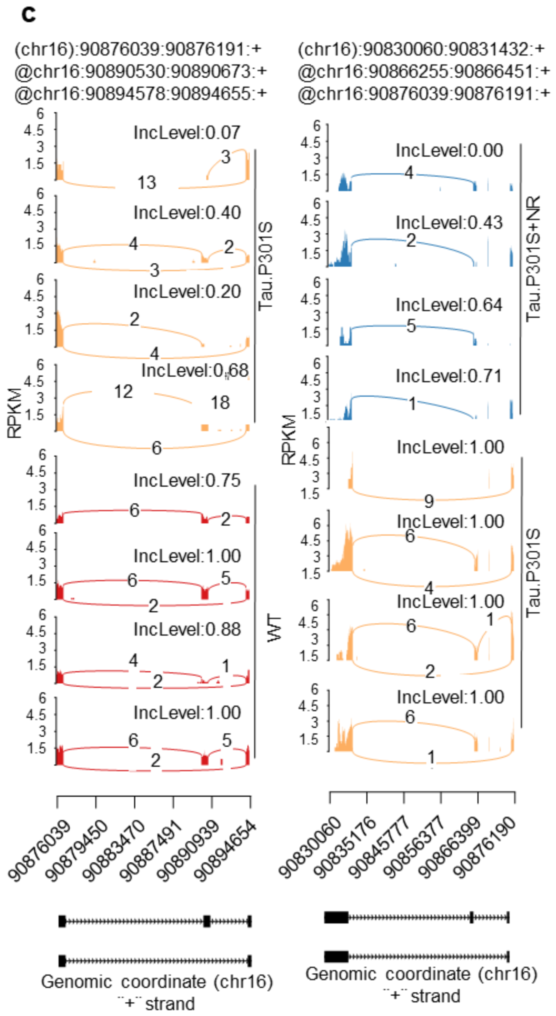


**Human cell line-based RNAseq data identify EVA1-C diverse sequences altered by NAD$^+$**

**a,** Heatmap showing different *Eva1-C* transcripts expression in human SH-SY5Y and 2N3R tau-overexpressing SH-SY5Y with- or without- NMN at 4 different doses. **b**, Summary of biotype of *EVA1-C* transcripts. **c**, Sashimi plot of EVA1-C splicing changes in WT (Veh.), hTau.P301S (Veh.), and hTau.P301S (NR) groups. Diagrams on the left show the read coverage of exons. Plots on the right show the incLevel values that occurred in two paired tissues. The AS model of this region is represented in the panels underneath. Each curve indicates the numbers of splicing sites and the number in the curve suggests the amount of RNA-seq reads in this region.

**Fig. S4.**

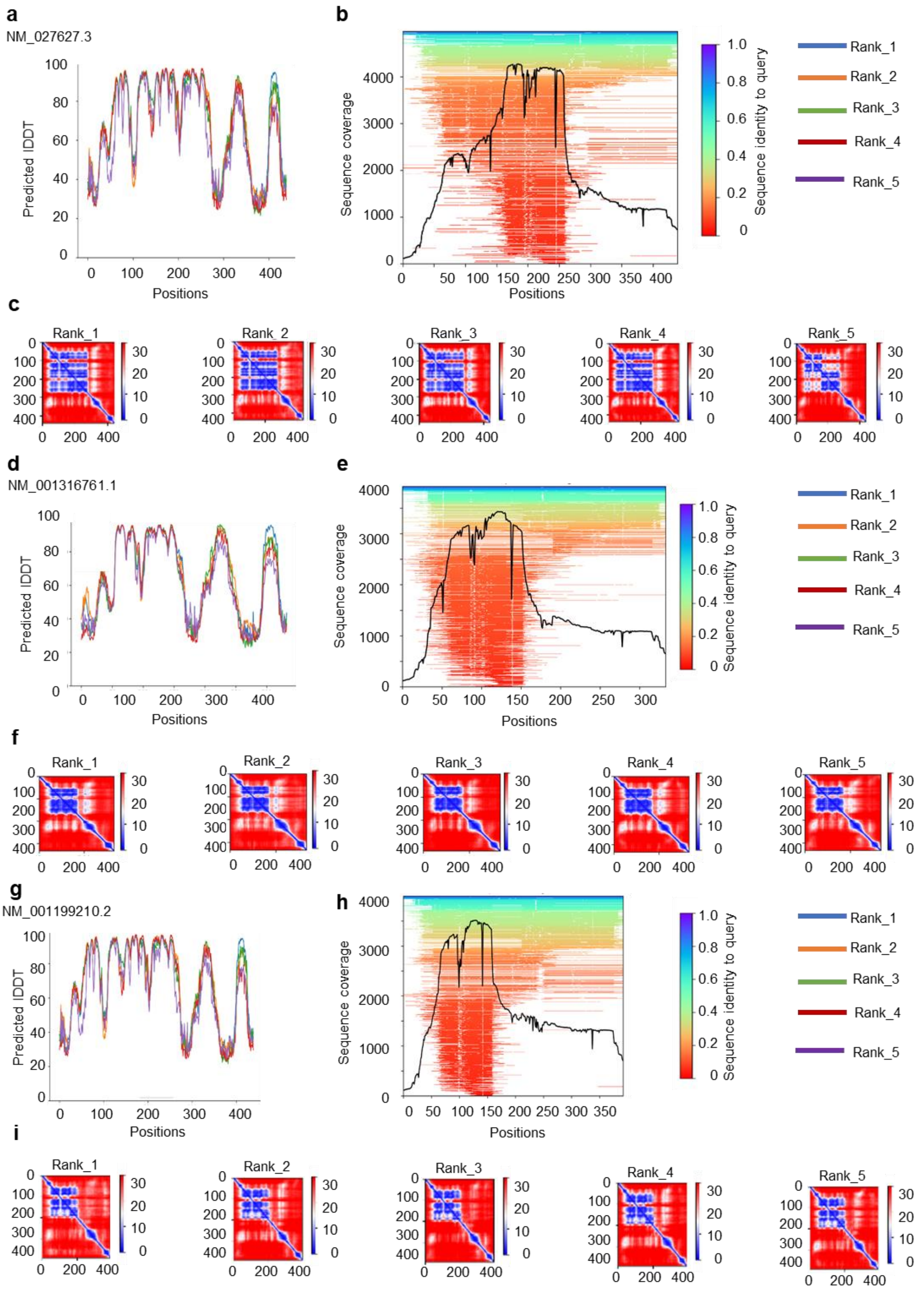
a
NM_027627.3
Predicted lDDT
Positions
b
Sequence coverage
Positions
Sequence identity to query
Rank_1
Rank_2
Rank_3
Rank_4
Rank_5
c
d
NM_001316761.1
e
f
g
NM_001199210.2
h
i

**Quality assessment of EVA1-C protein conformation prediction**

**a, d**, **g,** l*eft* the IDDT plot which is a local superposition-free score associated with model confidence, while a higher IDDT score correlates with better performance of the model at that location. **b, e**, **h,** *right* the plot for the number of sequences per position. The criteria are: higher than 30 sequences per position; the three isoforms have over 30 sequences per position and 93% of the sequence has over 100 reference sequences. **c, f, i,** the plot for predicted alignment error. This metric is applied to assess how confident the model is about the interface. The lower the score, the better.

**Fig. S5.**

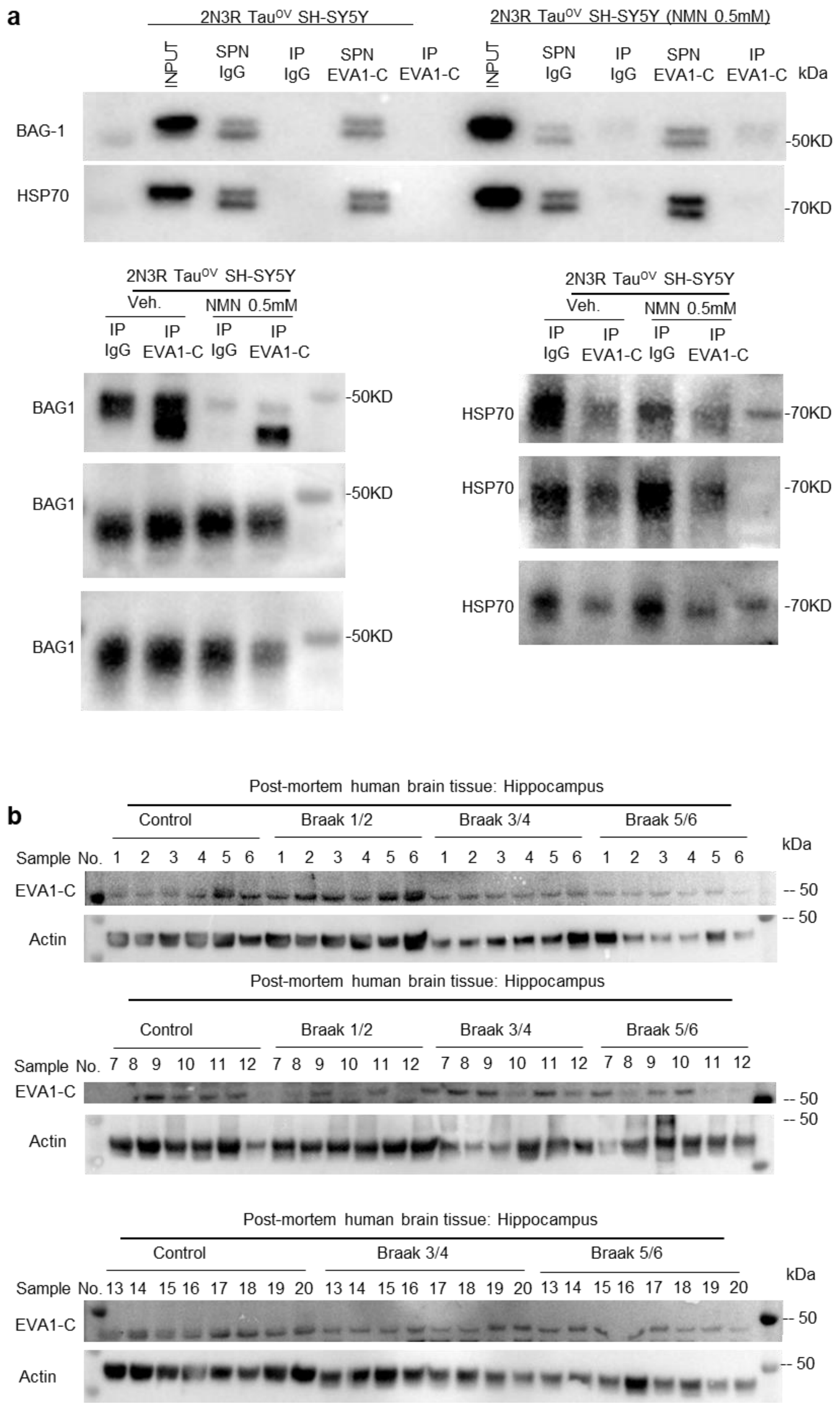
a
2N3R Tau^OV SH-SY5Y
2N3R Tau^OV SH-SY5Y (NMN 0.5mM)
INPUT
SPN IgG
IP IgG
SPN EVA1-C
IP EVA1-C
kDa
BAG-1
HSP70
-50KD
-70KD
Veh.
NMN 0.5mM
BAG1
-50KD
HSP70
-70KD
b
Post-mortem human brain tissue: Hippocampus
Control
Braak 1/2
Braak 3/4
Braak 5/6
Sample No. 1 2 3 4 5 6 1 2 3 4 5 6 1 2 3 4 5 6 1 2 3 4 5 6
EVA1-C
Actin
-- 50
Sample No. 7 8 9 10 11 12 7 8 9 10 11 12 7 8 9 10 11 12 7 8 9 10 11 12
Sample No. 13 14 15 16 17 18 19 20 13 14 15 16 17 18 19 20 13 14 15 16 17 18 19 20

**The $NAD^+$-EVA1-C axis regulates BAG-1 and HSP70 binding in human cells, as well as reduced EVA1-C protein in postmortem tissues from AD patients**

**a**, Co-immunoprecipitation (Co-IP) assays showing interactions of EVA1-C with HSP70 and BAG-1. **b**, Western blot data showing EVA1-C protein expression in hippocampal brain tissues from AD patients with different Braak stages and age-matched healthy control. Cognitive normal samples, n = 20; Braak 1/2, n = 12; Braak 3/4, n = 20; Braak 4/5, n = 20.

**Table S4.**

| *C. elegans* gene name | Mamalian protein homologue | Name | Function |
|---|---|---|---|
| *uaf-1* | U2AF35 | U2 auxiliary factor small subunit | Core spliceosomal factor |
| *uaf-2* | U2AF65 | U2 auxiliary factor large subunit | Core spliceosomal factor |
| *sfa-1* | SF1/BBP | Splicing factor 1, branch point binding protein | Core spliceosomal factor |
| *repo-1* | SF3A2 | Splicing factor 3a subunit 2 (66kDa) | Core spliceosomal factor |
| *snr-1* | SNRPD3 | Small nuclear ribonucleoprotein Sm D3 | Core spliceosomal factor |
| *snr-2* | SNRPB | Small nuclear ribonucleoprotein Sm B | Core spliceosomal factor |
| *rsp-2* | SRSF5, SRp40 | Serine/Arginine-rich splicing factor 5 | Extrinsic non spliceosomal RNA binding protein |
| *rsp-3* | SRSF1, SF2/ASF | Serine/Arginine-rich splicing factor 1 | Extrinsic non spliceosomal RNA binding protein |
| *hrp-1* | hnRNP A1 | Heterogeneous nuclear ribonucleoprotein A1 | Extrinsic non spliceosomal RNA binding protein |
| *hrp-2* | hnRNP R | Heterogeneous nuclear ribonucleoprotein R | Extrinsic non spliceosomal RNA binding protein |
| *hrpf-1* | hnRNP F/H | Heterogeneous nuclear ribonucleoprotein F/H | Extrinsic non spliceosomal RNA binding protein |
| *prp-8* | PRPF8 | Pre-mRNA processing splicing factor 8 | Core spliceosomal factor |
| *phi-9* | NHP2L1 | NHP2-like protein 1 | RNA binding protein |
| *smu-1* | SMU1 | WD40 repeat-containing protein SMU1, fSAP57 | RNA binding protein |
| *sym-2* | hnRNP F/H | Heterogeneous nuclear ribonucleoprotein F/H | Extrinsic non spliceosomal RNA binding protein |

**A summary of splicing factors in *C. elegans* and their mammalian homologues**

**Table S18.**

| Groups | Median lifespan (day) | Mean ± S.E.M. (day) | Statistics ($p$ values) |
| --- | --- | --- | --- |
| WT (Veh.) | 15.09 | 15.09 ± 0.5589 | |
| hTau[P301L] (Veh.) | 11.40 | 11.40 ± 0.3627 | ***p* < 0.0001 vs WT (Veh.)** |
| hTau[P301L] (NR) | 13.32 | 13.32 ± 0.4208 | ***p* = 0.0015** vs hTau[P301L] (Veh.) ** |
| hTau[P301L];*eva1-c(RNAi)*$^{n\text{-}sid\text{-}1OV}$ | 11.78 | 11.78 ± 0.3634 | *p* = 0.6427 vs hTau[P301L] (Veh.) |
| hTau[P301L];*eva1-c (RNAi)*$^{n\text{-}sid\text{-}1OV}$ (NR) | 11.36 | 11.36 ± 0.2618 | *p* = 0.4260 vs hTau[P301L] (Veh.) |

**A summary of the *C. elegans* lifespan data.**

**Table S20.**

| Samples | | Age | Sex | PMD (h) | Allele | Clinical Dignosis | Tauopathy | Exp. |
|---|---|---|---|---|---|---|---|---|
| **No.** | **ID** | | | | | | | |
| **1** | A393/19 | 92 | F | 63 | APOE 3/3 | use as control | Tau Braak stage 2 | W |
| **2** | A078/17 | 98 | F | 76 | APOE 3/3 | use as control | Tau Braak stage 2 | W |
| **3** | A237/16 | 80 | M | 58 | APOE 3/3 | use as control | | W, I |
| **4** | A066/16 | 95 | M | 72.5 | APOE 2/3 | use as control | | W |
| **5** | A007/15 | 74 | F | 66 | APOE 2/3 | use as control | Tau Braak stage 2 | W, I |
| **6** | A319/14 | 90 | F | 44 | APOE 3/3 | use as control | Tau Braak stage 2 | W |
| **13** | A302/18 | 90 | F | 30 | APOE 2/3 | use as control | | W |
| **14** | A049/18 | 75 | F | | APOE 3/3 | use as control | Tau Braak stage 1 | W |
| **15** | A226/17 | 90 | F | 48 | APOE2/3 | use as control | Tau Braak stage 1 | W |
| **16** | A382/16 | 87 | M | 48 | APOE 3/4 | use as control | | W |
| **17** | A297/16 | 81 | M | 10 | APOE 3/3 | use as control | Tau Braak stage 1 | W |
| **18** | A473/15 | 74 | M | 72 | APOE3/4 | use as control | | W |
| **19** | A242/15 | 82 | M | 26 | APOE3/4 | use as control | Tau Braak stage 2 | W |
| **20** | A234/15 | 80 | M | 34 | APOE3/4 | use as control | Tau Braak stage 1 | W |
| **1** | A142/16 | 83 | M | 48 | APOE 2/3 | Braak 1-2 | Tau Braak stage 2 | W, I |
| **2** | A046/14 | 74 | F | 72 | APOE 3/3 | Braak 1-2 | | W, I |
| **3** | A354/16 | 92 | F | 12 | APOE3/3 | Braak 1-2 | Tau Braak stage 2 | W |
| **4** | A073/05 | 93 | M | ~33.00 | APOE3/3 | Braak 1-2 | Tau Braak stage 2 | W |
| **5** | A187/06 | 71 | F | 48 | APOE3/4 | Braak 1-2 | Tau Braak stage 1 | W |
| **6** | A067/09 | 92 | F | 19.5 | APOE3/3 | Braak 1-2 | Tau Braak stage 3 | W |
| **1** | A046/13 | 85 | M | 54 | APOE 3/3 | Braak 3-4 | Tau Braak stage 3-4 | W, I |
| **2** | A097/13 | 82 | M | 28 | APOE4/4 | Braak 3-4 | Tau Braak stage 4 | W, I |
| **3** | A097/15 | 90 | F | 43 | APOE 3/3 | Braak 3-4 | Tau Braak stage 4 | W |
| **4** | A266/15 | 90 | F | 82.5 | APOE 3/3 | Braak 3-4 | Tau Braak stage 4 | W |
| **5** | A065/16 | 91 | M | 48 | APOE 2/4 | Braak 3-4 | Tau Braak stage 4 | W |
| **6** | A444/18 | 95 | M | 61 | APOE 2/3 | Braak 3-4 | Tau Braak stage 4 | W |
| **13** | A374/14 | 88 | M | 79 | APOE 3/4 | Braak 3-4 | Tau Braak stage 3-4 | W |
| **14** | A233/13 | 92 | M | 70 | APOE3/3 | Braak 3-4 | Tau Braak stage 4 | W |
| **15** | A078/13 | 86 | M | 52.5 | APOE3/4 | Braak 3-4 | Tau Braak stage 4 | W |
| **16** | A357/11 | 91 | M | 28 | APOE3/3 | Braak 3-4 | Tau Braak stage 4 | W |
| **17** | A362/18 | 92 | F | 55 | APOE 3/3 | Braak 3-4 | Tau Braak stage 4 | W |
| **18** | A418/17 | 94 | M | 62 | APOE 3/3 | Braak 3-4 | Tau Braak stage 4 | W |
| **19** | A381/16 | 84 | M | 86 | APOE 3/3 | Braak 3-4 | Tau Braak stage 4 | W |
| **20** | A305/09 | 81 | M | 13 | APOE 2/4 | Braak 3-4 | Tau Braak stage 3 | W |
| **1** | A221/13 | 89 | M | 26 | APOE3/4 | Braak 5-6 | Tau Braak stage 5 | W |
| **2** | A355/14 | 79 | F | 31 | APOE 3/3 | Braak 5-6 | Tau Braak stage 6 | W, I |
| **3** | A366/14 | 82 | F | 68 | APOE 3/4 | Braak 5-6 | Tau Braak stage 6 | W |

| | | | | | | | | |
|---|---|---|---|---|---|---|---|---|
| **4** | A377/14 | 85 | F | 79 | APOE 4/4 | Braak 5-6 | Tau Braak stage 6 | W |
| **5** | A395/14 | 92 | M | 36.5 | APOE 4/4 | Braak 5-6 | Tau Braak stage 5 | W, I |
| **6** | A156/15 | 86 | F | 43.5 | APOE 4/4 | Braak 5-6 | Tau Braak stage 6 | W |
| **13** | A258/16 | 67 | M | 39.5 | APOE 4/5 | Braak 5-6 | Tau Braak stage 6 | W |
| **14** | A166/16 | 89 | M | 21 | APOE 3/2 | Braak 5-6 | Tau Braak stage 6 | W |
| **15** | A087/16 | 89 | F | 38.5 | APOE 3/3 | Braak 5-6 | Tau Braak stage 6 | W, I |
| **17** | A163/15 | 76 | F | 4 | APOE4/4 | Braak 5-6 | Tau Braak stage 6 | W |
| **18** | A092/15 | 86 | F | 13 | APOE3/4 | Braak 5-6 | Tau Braak stage 6 | W, I |
| **19** | A342/14 | 84 | F | 27 | APOE 3/4 | Braak 5-6 | Tau Braak stage 6 | W |
| **20** | A118/20 | 87 | F | 44 | APOE 3/3 | Braak 5-6 | Tau Braak stage 6 | W |
| | A141/18 | 82 | F | 42 | APOE3/3 | Control | | I |
| | A030/19 | 98 | F | 9 | APOE3/3 | Control | Tau Braak stage 1 | I |
| | A364/11 | 96 | F | 33 | APOE3/4 | Braak 1-2 | Tau Braak stage 2 | I |
| | A344/16 | 77 | M | 96 | APOE3/3 | Braak 1-2 | Tau Braak stage 2 | I |
| | A232/16 | 95 | F | 47 | APOE3/4 | Braak 3-4 | Tau Braak stage 4 | I |
| | A084/16 | 86 | F | 55.5 | APOE3/4 | Braak 3-4 | Tau Braak stage 4 | I |
| | A259/15 | 92 | F | 21 | APOE4/4 | Braak 5-6 | Tau Braak stage 5 | I |

**Detailed information of individuals with their postmortem brain tissues used in this study [western blotting (W), and immunostaining (I)]**
